# Spatial multi-omics and single-cell transcriptomics uncover senescence-associated cellular programs during colon adenoma to cancer progression

**DOI:** 10.64898/2026.08.06.743310

**Authors:** Ming Yu, Yumo Xie, Lena Allen, Kelly Carter, Elizabeth Donato, Lingxin Cheng, Christina Kendziorski, Kristina A. Matkowskyj, Angelo M. DeMarzo, Jessica Hicks, Deepti Reddi, Evan W. Newell, Wei Sun, William M. Grady

## Abstract

Colorectal cancer develops through a normal-adenoma-carcinoma sequence, yet only 5-10% of adenomas progress to malignancy, and the cellular programs governing that sequence remain poorly defined. Here we generate a spatial multi-omics atlas of human colon adenomas, combining Visium CytAssist and protein co-detection across 24 nonadvanced and advanced tubular adenomas with single-cell resolution Xenium Prime 5K profiling of 101 patient-matched normal, adenoma, and carcinoma cores from 16 patients. Integrating whole-transcriptome and 31-plex protein data identifies nine spatial clusters and two dysplastic epithelial populations that co-express stemness, proliferation, and senescence programs. These programs occupy a shared, spatially confined epithelial niche that expands from adenoma to carcinoma. Spatial analysis revealed GDF15, a senescence-associated secretory factor, mediated the coupling between senescence and stemness in advanced adenomas, and that GDF15-high epithelium locally excludes CD8+ T cells in adenoma and, more broadly, in carcinoma. These findings position senescence as a spatially instructive rather than merely tumor-suppressive program during colorectal carcinogenesis and suggest GDF15 might be a potential candidate target for cancer prevention and interception in the colon.

**Statement of Significance:** Spatial multi-omics identifies spatially co-localized senescence and stemness programs in dysplastic epithelium during colorectal tumorigenesis and links GDF15-associated niches to CD8-positive T-cell exclusion, nominating GDF15 for colorectal cancer prevention and interception strategies.

**Conflicts of interests:** The authors declare the following competing interests: W.M.G.: Consulting for Guardant Health, Research support from Lucid Diagnostics; E.W.N.: Co-founder, advisor, and shareholder of ImmunoScape. All the other authors declare no competing interests.

## Introduction

Colorectal cancer (CRC) arises through a well-established normal-adenoma–carcinoma sequence, during which molecular changes in the normal colonic epithelium accumulate and mediate the histologic steps towards cancer formation. It is notable that only a minority (5–10%) of adenomas progress to cancer. The determinants that govern this divergent trajectory remain poorly defined. Genetic and epigenetic alterations in the neoplastic epithelium, including mutations in APC, KRAS, and TP53, have long been recognized as central drivers of adenoma-to-carcinoma progression^1,2^. More recently, single-cell transcriptomic and epigenomic profiling has revealed that a large fraction of polyp and cancer cells occupy a stem-like state, and that the transition from normal to malignancy follows a continuous epigenetic and transcriptional trajectory rather than a series of discrete molecular steps^3,4^. Recent lineage-tracing studies in mouse models of intestinal tumorigenesis have demonstrated that early colorectal lesions are often polyclonal in origin, comprising multiple independent cell lineages undergoing parallel clonal expansions, with extensive intercellular cooperation preceding a later polyclonal-to-monoclonal transition that marks disease advancement^5^. These findings collectively underscore that colon cancer initiation and progression is not simply a monoclonal, cell-autonomous process but is instead shaped by complex interactions between neoplastic epithelial states and the surrounding tissue microenvironment. A more integrated understanding of these early events is critical for developing effective risk stratification, prevention, and interception strategies.

Colorectal cancer is fundamentally a disease of aging, and advanced age remains one of the most significant risk factors for CRC. Cellular senescence, an irreversible state of cell-cycle arrest induced by oncogenic stress, genotoxic insult, or replicative exhaustion, has emerged as a convergence between hallmarks of aging and precancerous lesions^6,7^. Senescent cells acquire a senescence-associated secretory phenotype (SASP) that profoundly reshapes the tissue milieu through cytokine, chemokine, and growth factor release. Senescence is canonically considered a tumor-suppressive mechanism that limits the proliferation of damaged cells, however, persistent or spatially organized senescence within the precancer microenvironment may paradoxically establish permissive niches that enable immune evasion and tumor^8^. In established CRC, distinct senescent cancer cells have been identified that correlate with the spatial evolution toward more metastatic phenotypes at the invasive front^9^. Little is known about how senescence is spatially organized within adenomatous tissue, which cell populations harbor this program, and whether it coordinates with stemness and immune exclusion programs across the adenoma-to-carcinoma continuum.

To spatially profile the colon adenoma and test the hypothesis that senescence plays an important role in colon cancer initiation and progression, we generated a comprehensive spatial multi-omics atlas of human colon adenomas encompassing non-advanced and advanced lesions, using the Visium spatial transcriptomics platform with protein co-detection. To validate and extend these findings at single-cell resolution across the full disease spectrum, we constructed a tissue microarray comprising normal, adenoma, and carcinoma tissues and applied single-cell resolution spatial transcriptomics. Together, the resulting datasets enabled us to map the cellular composition, transcriptional state, and spatial organization of the adenoma microenvironment at an unprecedented single-cell resolution. Integration of these two complementary spatial modalities reveals that senescence and stemness, classically viewed as opposing cellular states, co-existed in dysplastic epithelial niches. These epithelial niches emerged in adenomas, expanded with lesion progression, and are spatially organized around GDF15-expressing cells that actively exclude CD8+ T cells from their immediate microenvironment. These findings redefine senescence in the precancerous colon as a spatially instructive program that coordinates epithelial plasticity and immune evasion, and identify candidate molecular targets, such as GDF15, for colorectal cancer prevention and interception.

## Results

### Multimodal spatial mapping delineates the dysplastic epithelium in tubular adenomas

To spatially map the colon adenomas, we performed Visium Spatial Gene Expression and Protein co-detection (Figure 1). Twenty-four conventional adenomas (AD, including tubular/ tubulovillous adenomas) were initially profiled (8 nonadvanced and 16 advanced). Two nonadvanced adenomas failed prespecified quality-control criteria, leaving 22 specimens for analysis (6 nonadvanced and 16 advanced) (Supplementary Table 1). We integrated the RNA and antibody-derived tags (ADTs) data to improve delineation of epithelial, stromal, and immune compartments (Supplemental Figure 1A and 1B). Single-spot Uniform Manifold Approximation and Projection (UMAP) visualization and clustering identified nine spatial clusters (C0-C8), each defined by distinct marker-gene and representative ADT signals (Figure 2A and 2B; Supplementary Figures 1C and 2). Using histologic annotation, we classified Visium spots as stroma, normal epithelium, or adenoma (Methods). All adenoma regions in this cohort were low-grade dysplasia (Figure 2C; Supplementary Figures 1D, 1E and Supplementary Figure 2).

**Figure 1.**
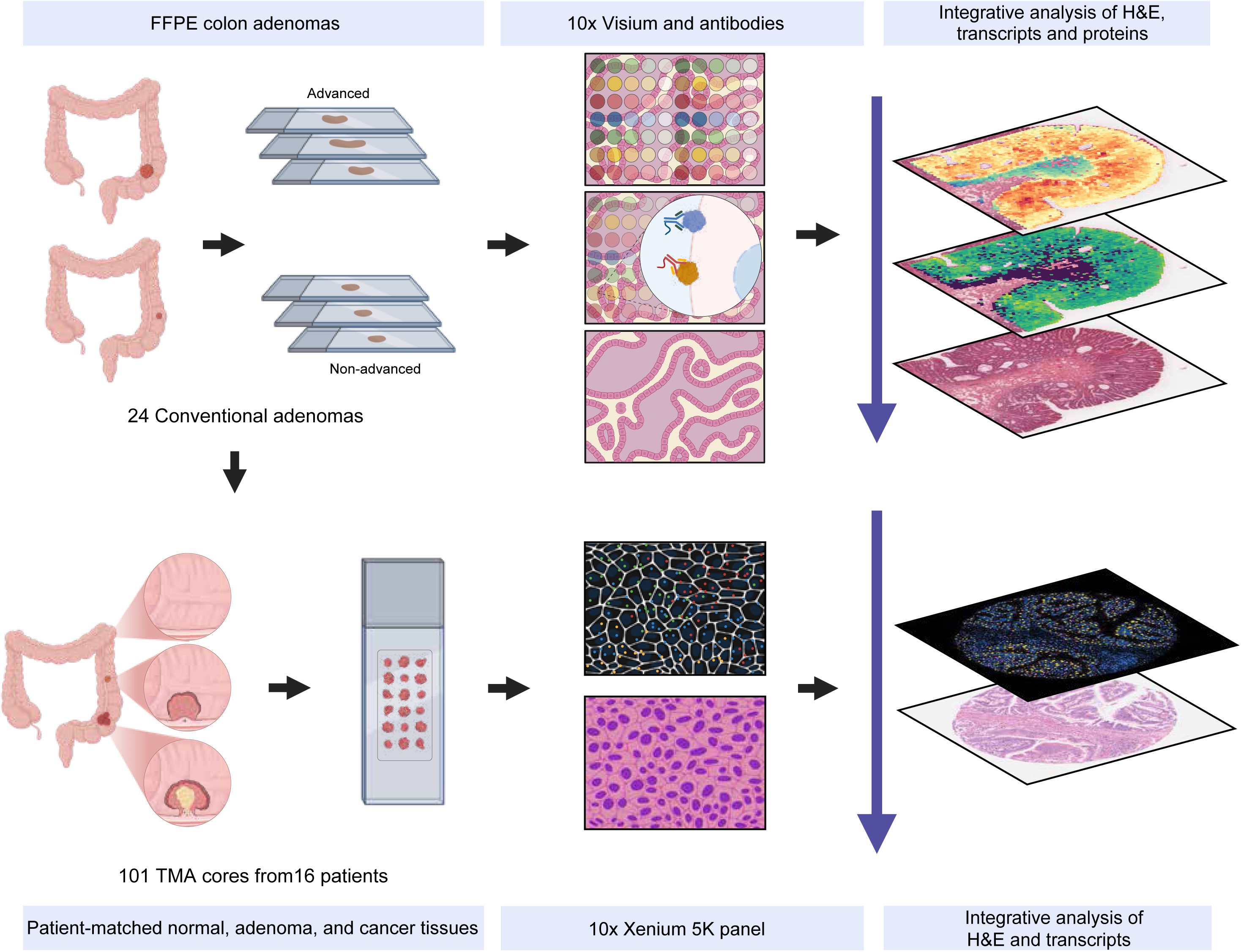
Schematic diagram depicting the workflow for integrated multimodal spatial profiling of colon adenomas at different points in the adenoma to adenocarcinoma progression sequence. We employed a trimodal integration approach combining H&E-based pathology with simultaneous RNA and protein detection at both tissue-level and subcellular resolution. The upper panel shows 24 FFPE conventional adenomas, stratified by size and histology (advanced and non-advanced), were profiled using Visium CytAssist Spatial Gene and Protein Expression, which co-detects whole transcriptome gene expression and DNA-barcoded antibody-based protein markers on the same tissue section. The lower panel shows 101 tissue microarray (TMA) cores from 16 patients, comprising patient-matched normal, adenoma, and cancer tissues, which were analyzed using Xenium in situ transcriptomics and post-Xenium H&E. These platforms enabled integrative analysis of tissue morphology (H&E), spatially resolved transcript profiles, and protein localization patterns, providing a comprehensive multimodal characterization of cellular heterogeneity and the tumor microenvironment across the adenoma-carcinoma sequence.

**Figure 2.**
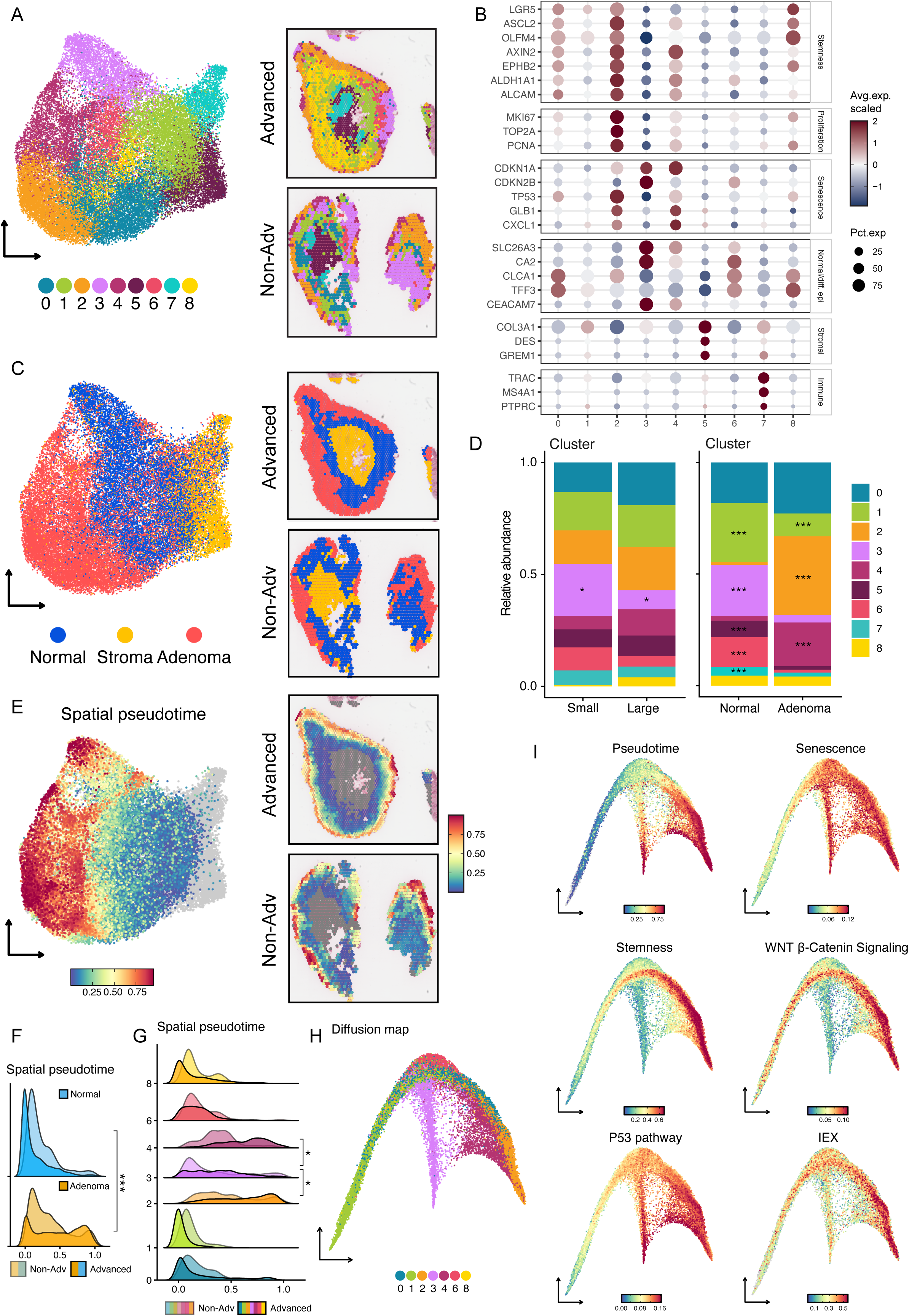
Integrated multimodal annotation of colon adenoma Visium spatial transcriptomics. Visium CytAssist Spatial Gene and Protein Expression data were integrated using weighted nearest neighbor (WNN) analysis to combine RNA and DNA-barcoded antibody (ADT) information for comprehensive cellular characterization. (A) Left: UMAP visualization of integrated clustering across all samples using WNN analysis. Nine distinct clusters (0-8) were identified based on the weighted combination of both modalities. Right: Representative spatial maps showing clusters distribution in advanced (top) and nonadvanced (bottom) adenomas. (B) Dot plot showing the representative markers with functional annotation for each cluster. (C) Left: UMAP visualization of pathology-based annotation for all spots, displaying tissue compartments including normal epithelium, stroma, and adenoma. Right: Representative spatial maps showing pathology annotation in advanced (top) and nonadvanced (bottom) adenomas. (D) Distribution of integrated clusters between nonadvanced and advanced adenomas (Left) and between normal epithelium and adenoma regions (right). (E) Left: UMAP visualization of the spatial pseudotime for all spots. Right: Representative spatial maps showing the spatial pseudotime in advanced (top) and nonadvanced (bottom) adenomas. (F) Ridgeline plot showing the distribution of spatial pseudotime in normal epithelium and adenoma from nonadvanced and advanced samples. (G) Ridgeline plot showing the distribution of spatial pseudotime in spatial clusters from nonadvanced and advanced samples (H) Diffusion map visualization of the epithelial compartment, immune and stromal compartment were excluded. DC1 and DC3 was chosen for visualization. (I) Visualization of the level of different scores in the diffusion map space for the epithelial compartment.

Integrating the ADT signals along with the RNA marker gene, we classified C5 as stromal and C7 as immune, with the remaining seven clusters as distinct epithelial populations (Supplementary Figure 3A). The epithelial cluster-specific marker genes indicative of different cellular programs (Figure 2B). Stemness and cancer stem-like genes were most highly expressed in C0, C1 and C2, while proliferation markers were upregulated in C2. High expression in senescence-related genes^10^ were found C2, C3 and C4, and markers of mature differentiated epithelium were mainly restricted to C3 and C6. Further immunohistochemistry staining of CDKN1A (p21), CDKN2A (p16), and MKI67 (Ki67) showed high consistency with Visium spatial RNA expression, supporting these patterns (Supplementary Figure 3B).

This cluster-specific marker distribution informed our functional classification of different clusters. Of note, C1 shows moderate expression of LGR5 and OLFM4, but no proliferation or senescence activity, corresponding roughly to the crypt base epithelium. C2 was unique in co-expressing all three dysplastic programs simultaneously: stemness, proliferation, and senescence. C4 carried a distinct senescence signature dominated by CDKN1A without accompanying proliferation, consistent with a growth-arrested, oncogene-induced senescent state^11^. C3 and C6 represent differentiated normal epithelium, with C3 showing the highest expression of CDKN2B. Sample-based proportion testing reveals a significant reduction of C3 in advanced compared with nonadvanced adenomas (adj. P < 0.05). More interestingly, within-sample comparisons revealed marked expansion of C2 and C4 along with depletion of C1, C3 and C6 in adenoma regions relative to adjacent normal epithelium (Figure 2D), further supporting the classification of C2 and C4 as the principal dysplastic clusters.

To position these clusters along a developmental axis, we performed spatial pseudotime analysis on all high-confidence epithelial spots, designating C1 as the root within each sample (Figure 2E). We observed a significantly higher spatial pseudotime in the adenoma spots compared to normal spots within the same sample (P < 0.001; Figure 2F, sample-level paired Wilcoxon test). At the cluster level, C2 and C4 advanced further than C3, consistent with their dysplastic classification (Figure 2G, C2 vs. C3, P = 0.03; C4 vs. C3, P = 0.01; sample-level paired Wilcoxon test). To reconstruct a broader developmental trajectory, we then performed unsupervised diffusion map analysis in the epithelial clusters. This placed C1 at one extreme pole while C2 and C3 at two separate poles, validating C1 as the root of the trajectory and a bifurcation between the dysplastic (C2) and differentiated branches (C3) (Figure 2H).

Gene set enrichment analysis, aggregated across four scoring algorithms by Robust Rank Aggregation, showed that both dysplastic clusters, C2 and C4, were enriched for stemness, TGF-β signaling, and immune exclusion signature (IEX). C2 was further distinguished by Myc target activation, Wnt/β-catenin signaling, and G2/M checkpoint activity, alongside reduced inflammatory signaling (Figure 2I, Supplementary Figure 3C and 3D)^12^. Cell cycle scoring confirmed the highest G2/M occupancy in C2 and predominant G1 arrest in C3, the differentiad cluster (Supplementary Figure 3E and 3F). Senescence scoring by SenePy^13^, which computes a background-subtracted senescence score, showed a gradual increase along the trajectory, with C4 reaching the highest values. Although the absolute differences in median scores between clusters were modest (C2 = 0.080, C3 = 0.074, C4 = 0.089), C4 scores were consistently higher across adenoma samples (C4 vs. C3, adj P = 0.005; C4 vs. C2, adj P = 0.005; Supplementary Figure 3G). Together, these results spatially delineated the dysplastic and normal tissue within the colon adenoma.

### Co-existence of senescence, stemness, and immune suppression in the adenoma microenvironment

While senescence and stemness have been viewed as two opposing states, recent evidence suggests senescence can paradoxically induce stemness in certain cancer contexts^14,15^. When we projected both scores onto the integrated UMAP, we observed senescence and stemness signals overlapped spatially (Figure 3A, Supplementary Figure 4A). To independently assess the stem-like character of these regions, we estimated developmental potential using CytoTRACE, for which higher scores indicate a less differentiated state. CytoTRACE scores concorded with the stemness signature, supporting the presence of less differentiated epithelial states within the senescence-stemness-associated regions (Supplementary Figure 4B).

**Figure 3.**
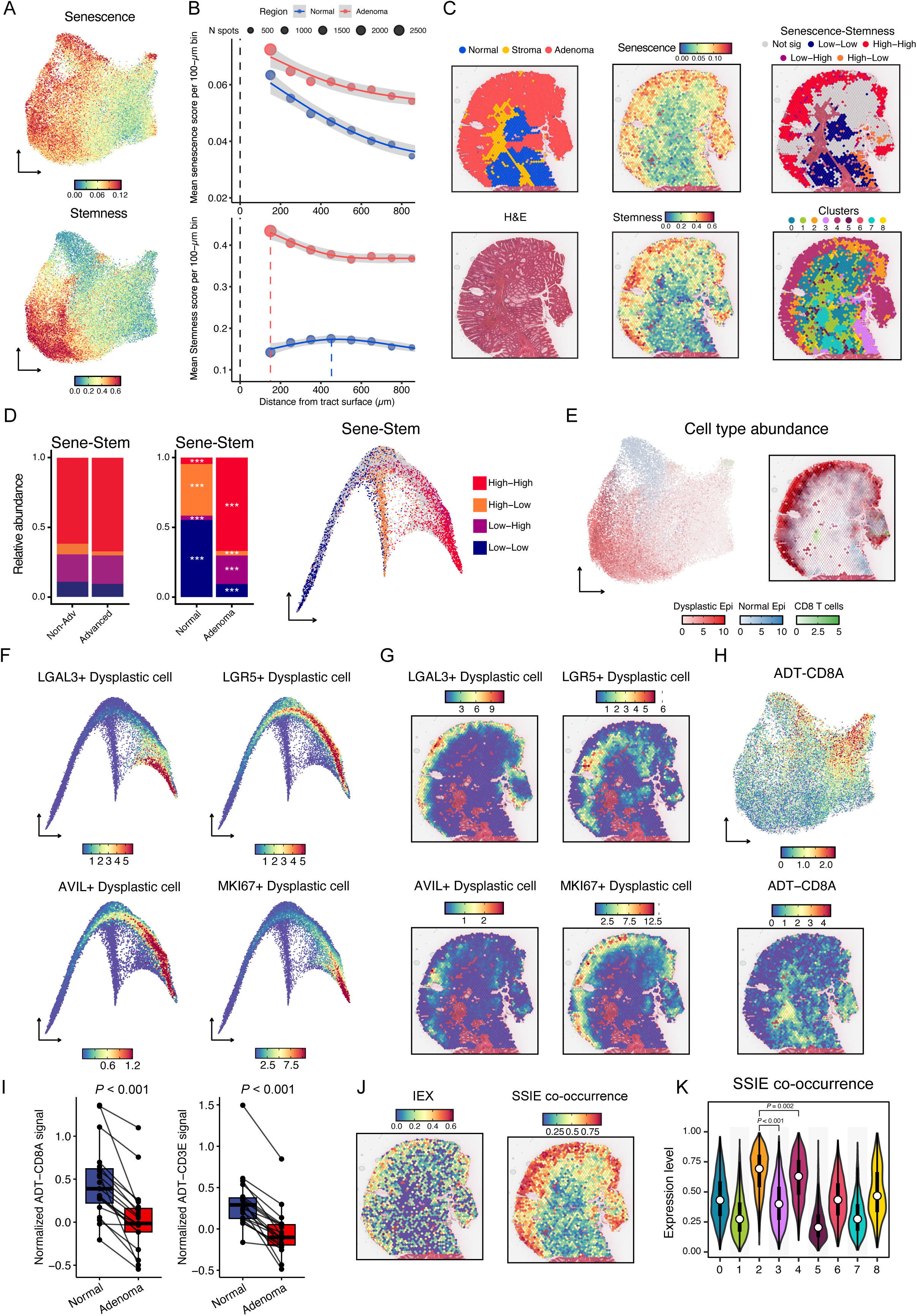
Cellular senescence and stemness characterization reveal spatial heterogeneity and association with adenoma progression. (A) UMAP visualization of the senescence (Top, by SenePy Universal hub) and Stemness (Bottom, by stemness gene signature) for all spots. (B) Non-stromal Visium spots were grouped into 100-µm bins according to their distance from the manually annotated luminal surface. Points show mean scores within 100-µm distance bins, with point size proportional to the number of spots; lines show generalized additive model-smoothed trends and shaded bands denote 95% confidence intervals. (C) Representative spatial maps of a advanced adenoma displaying pathological annotations, H&E staining, senescence, stemness, local bivariate Moran’s I classes and spatial cluster assignments. (D) Proportional distribution of local bivariate Moran’s I classes examining the spatial co-localization of senescence and stemness scores, stratified by advancement (left) and pathological annotation (middle), and the classes distribution in the diffusion map space (right). For comparison between advanced and non-advanced adenomas, only pathological adenoma regions were included. (E) Top: UMAP visualization of the estimated cell numbers for each spot by Cell2location deconvolution. The number of dysplastic epithelial, normal epithelial and CD8 T cells are illustrated by different colors. Bottom: The representative spatial map of a advanced adenoma. (F) Diffusion map visualization of four different functional subtypes of dysplastic epithelium. (G) Representative spatial maps of an advanced adenoma showing the spatial overlapping of these four dysplastic epithelial subtypes. (H) Top: UMAP visualization of the CD8A ADT intensity. Bottom: The representative spatial map of an advanced adenoma. (I) Box plots comparing normalized ADT-CD8A and ADT-CD3E signals between normal and adenoma regions. Lines connect matched regions from the same sample (n = 19); paired Wilcoxon signed-rank tests were used. (J) Representative spatial maps showing the Immune exclusion signature (IEX) and the Senescence–Stemness–Immune Exclusion Co-occurrence Score (SSIE). (K) Violin plots showing representative SSIE Co-occurrence Score across clusters. Statistical tests were performed at sample-level (n = 22) using paired Wilcoxon signed-rank tests. *, P < 0.05; ***, P < 0.001.

To examine whether these programs are organized along the crypt, we binned spots by distance from the luminal surface and plotted mean scores per 100-µm interval (Figure 3B). Senescence scores were highest near the luminal surface in both normal and adenoma regions and declined with depth, indicating that surface-proximal senescence is a shared feature of the colonic epithelium. Stemness followed a strikingly different pattern: in adenoma, scores were markedly elevated immediately beneath the luminal surface and fell sharply with increasing depth, whereas in normal epithelium stemness peaked near the base of the crypt, consistent with the known depth of a typical colon crypt (∼400–600 µm). This surface-confined emergence of high stemness was therefore adenoma-specific, and points to a spatial zone just below the luminal surface where both programs are simultaneously elevated in dysplastic tissue.

To quantify this spatial colocalization, we performed bivariate local Moran’s I analysis on senescence and stemness scores across all epithelial spots. We found “Sen^High^-Stem^High^” spots, where both scores were locally elevated, were spatially aligned with the dysplastic C2 and C4 clusters, while “Sen^Low^-Stem^Low^” spots dominated normal and stromal areas (Figure 3C). Within matched samples, adenoma regions were significantly enriched for Sen^High^-Stem^High^ spatial colocalizations compared with adjacent normal regions, accounting for the distinct overall distribution of bivariate stemness–senescence association classes (PERMANOVA P = 0.001; dispersion test P = 0.076; paired post-hoc Wilcoxon for High-High, adjusted P < 0.001), though the increase of Sen^High^-Stem^High^ in the adenoma region from nonadvanced to advanced samples is not statistically significant. Interestingly, on the diffusion map, the Sen^High^-Stem^High^ spots enriched at one extreme while Sen^High^-Stem^Low^ spots, representing areas of high senescence without adjacent stemness, enriched at a different branch, suggesting the separation of these two biological programs (Figure 3D).

Since each Visium spot can contain multiple cell types, we next performed spatial deconvolution to resolve the cellular contributors to the senescence–stemness colocalized niche. We applied Cell2location using a CRC single-cell atlas^16^ restricted to colonoscopy-derived AD (Supplementary Figure 4C). The original atlas cell labels were used as broad reference annotations, and the epithelial/tumor-labeled compartment from AD samples was further subclustered. In this data, both senescence and stemness programs were primarily enriched within the epithelial compartment (Supplementary Figure 4D). Further epithelial subclustering suggested that these two programs were preferentially enriched in distinct adenoma-derived epithelial states, rather than uniformly co-activated within the same one (Supplementary Figure 4E–H). Using this reference, spatial deconvolution indicated that Visium spots within dysplastic regions harbor multiple adenoma-derived epithelial subtypes, revealing pronounced epithelial heterogeneity in senescence–stemness colocalized areas (Figure 3E, Supplementary Figure 5). Histologically defined dysplastic regions with high senescence–stemness scores were predicted to contain mixtures of LGR5□ dysplastic cells with strong stemness programs and LGAL3□/AVIL□ dysplastic cells with higher senescence-associated activity (Figure 3F–G, Supplementary Figure 4E–H). These results suggest that the co-existence of senescence and stemness within the adenoma niche arises from the spatial co-localization of distinct cell states.

We further validated these findings at single-cell resolution by analyzing publicly available scRNA-seq data from two colon adenoma cohorts^4^. Consistently, this single-cell analysis supported the presence of distinct dysplastic epithelial subpopulations with separate activation of senescence and stemness programs within the adenoma niche (Supplemental Figure 6A-J); and the dysplastic epithelial cell in the adenoma is the major source of senescence signal (Supplemental Figure 6K).

We also mapped CD8+ T cell abundance by transcriptomic deconvolution and orthogonal ADT-based protein detection and found low CD8A signals in regions of highest dysplastic cell abundance (Figures 3E, 3H 3I, Supplementary Figures 4I). To directly quantify the concurrent enrichment of senescence, stemness, and immune exclusion, we calculated the senescence–stemness–immune exclusion (SSIE) co-occurrence score (Methods). High SSIE scores localized predominantly to dysplastic regions and were highest in C2, which occupies the terminal end of the trajectory (Figure 1H), supporting the spatial co-occurrence of senescence, stemness, and an immune-excluded microenvironment within this dysplastic state (Figures 3J, 3K and Supplementary Figure 4J). Together, these data define a spatially organized niche in which senescence and stemness programs co-localize within distinct dysplastic epithelial subpopulations and are associated with immune exclusion.

### Identification of GDF15 as a molecular mediator of the dysplastic stem-senescence niche

Having demonstrated that dysplastic C2 and C4 niches are defined by senescence–stemness colocalization and immune exclusion, we next assessed the senescence-associated secretory profiles of these niches to determine if these factors might mediate the stemness and immune exclusion states we have characterized. Because senescence is context-dependent^17^, and our data indicate that senescence and stemness signals arise from distinct subpopulations, we reasoned that paracrine signaling, rather than cell-autonomous signaling, was likely to mediate the spatial co-occurrence of these two programs. We therefore integrated three spatial DEG contrasts: (1) advanced versus nonadvanced adenomas; (2) stemness-high versus stemness-low dysplastic spots; (3) senescence-high adenoma versus senescence-high normal epithelial regions, to nominate genes associated with senescence in dysplastic rather than in normal epithelium.

The intersection of these contrasts identified six secreted candidate genes: GDF15, MMP7, VWA2, COL9A2, CNTN3 and THNSL2 (Figure 4A). Several of the additional candidates, including MMP7 and VWA2, have previously been implicated in colorectal neoplasia or senescence-associated programs, providing biological support for this spatially derived candidate set^9,18^. We focused subsequent analyses on GDF15, the most highly expressed candidate, a prominent stress-induced cytokine and a potent component of the senescence-associated secretory phenotype (SASP) with reported roles in promoting tumor cell stemness, tissue remodeling, and immune exclusion^19^. To account for the nesting of spots, we confirmed the association of GDF15 with adenoma progression, stemness-high dysplastic regions, and senescence-high dysplastic epithelium using sample-level expression summaries (Supplementary Figure 7A).

**Figure 4.**
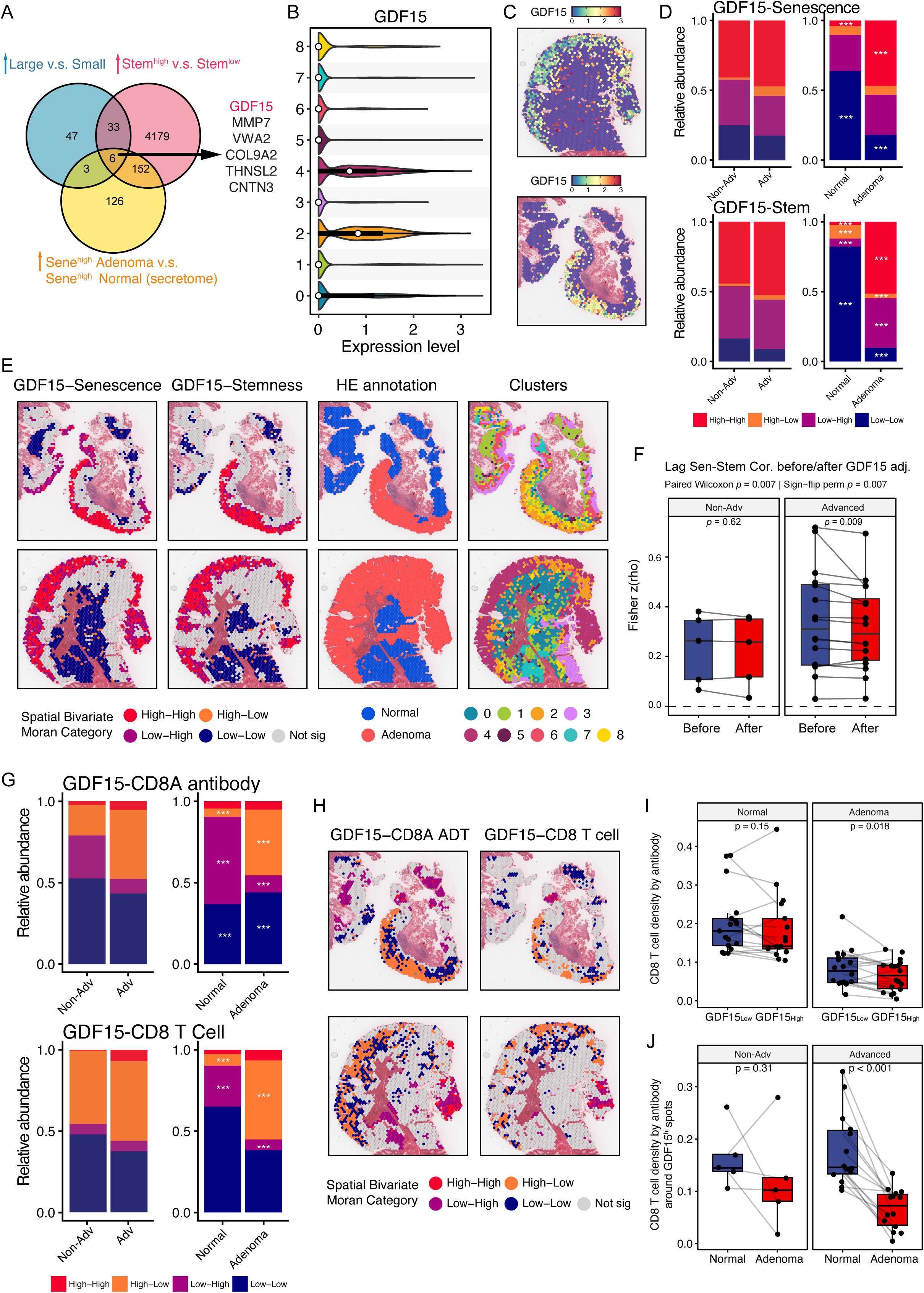
Spatial profiling identifies GDF15 as a coordinator of senescence, stemness, and CD8+ T cell exclusion in adenomas (A) Venn diagram illustrating the intersection of three comparisons: the C2 and C4 spots from advanced samples versus those from nonadvanced samples (Blue); Stemness^high^ versus Stemness^low^ spots in C2 and C4 (Red); SenePy^high^ spots from adenoma region versus those from normal region, featuring the secretome (Yellow). (B) Violin plots showing the expression GDF15 across clusters. (C) Representative spatial maps of GDF15 expression in two advanced adenomas (D) Stacked bar plots showing the proportional distribution of local bivariate Moran’s I classes, with GDF15-Senescence (Top) and GDF15-Stemness (Bottom). Colors denote Moran’s I classes. For comparison between advanced and non-advanced adenomas, only pathological adenoma regions were included. (E) Representative spatial maps of two advanced adenomas showing the local bivariate Moran’s I classes, H&E annotation and spatial clusters. (F) Box plot showing spatially smoothed senescence-stemness association before and after GDF15 adjustment in nonadvanced (n = 5) and advanced adenomas (n = 16). (G) Stacked bar plots showing the proportional distribution of local bivariate Moran’s I classes, with GDF15-CD8A antibody intensity (Top) and GDF15-CD8 T cell number by deconvolution (Bottom). For comparison between advanced and non-advanced adenomas, only pathological adenoma regions were included. (H) Representative spatial maps of two advanced adenomas showing the local bivariate Moran’s I classes. Colors denote Moran’s I classes. (I) Box plot showing the CD8 T cell density by ADT around the GDF15^high^ and GDF15^low^ spots in the normal and adenoma region. (J) Box plot showing the CD8 T cell density by ADT around the GDF15^high^ spots in the normal and adenoma region from nonadvanced (n = 5) and advanced adenomas (n = 14). ***, P < 0.001.

Mapping of GDF15 expression across the nine clusters indicated that GDF15 is highly upregulated in C2 and C4, confined to the dysplastic regions of adenoma tissue while absent from normal and stromal areas (Figure 4B, representative advanced adenomas in Figure 4C, Supplementary Figure 7B). Bivariate local Moran’s I analysis revealed that the spatial composition of the GDF15^High^-Sen^High^ category increased significantly in adenoma regions compared to the normal, concomitant with a significant decreased level in the GDF15^Low^-Sen^Low^ region. Similar patterns between GDF15 and stemness were observed, with additional significant increase in the GDF15^Low^-Stem^High^ region, as well as a significant decrease in the GDF15^High^-Stem^Low^ region in adenoma compared to normal (Figure 4D, post-hoc Wilcoxon, adj. P < 0.001 for all categories). Spatial maps confirmed that these GDF15^High^-Sen^High^ zones overlapped with the dysplastic C2 and C4 clusters (Figure 4E, Supplementary Figure 7G). Publicly available scRNA-seq data also validated that the GDF15 was predominantly expressed in adenoma cells and was highest in clusters with high scores in both stemness and senescence (Supplementary Figure 7C-F)^4^.

To further assess whether GDF15 may contribute to the stem-senescence co-localization, we calculated a spatial lag correlation between senescence at each spot and stemness in its surrounding neighborhood, before and after adjusting both programs for GDF15 expression (Methods). In nonadvanced adenomas, GDF15 adjustment had no significant effect on the senescence-stemness association (paired Wilcoxon *P* = 0.62). In advanced adenomas, however, controlling for GDF15 led to a modest but significant reduction in the spatial lag correlation between senescence and stemness (paired Wilcoxon P = 0.009, sign-flip permutation P = 0.007; Figure 4F), consistent with GDF15 contributing to their spatial coupling in advanced lesions. In addition, spatial lag correlations between GDF15 and either senescence or stemness were modest but consistently positive and higher in advanced than in non-advanced adenomas (pooled median Fisher z: GDF15–Sen, 0.05 in non-advanced vs 0.14 in advanced; GDF15–stemness, 0.15 in non-advanced vs 0.19 in advanced), indicating that GDF15 is spatially associated with both programs. Together with the enrichment of GDF15^High^–Sen^High^ and GDF15^High^–Stem^High^ categories in adenoma regions (Figure 4D), these findings suggest that GDF15 participates in the shared senescence–stemness niche rather than acting as an arbitrary covariate.

We then examined the spatial relationship between GDF15 expression and CD8+ T cell localization using bivariate Moran’s I analysis (Figure 4G, 4H). To ensure robust quantification, CD8+ T cell localization was evaluated orthogonally using both transcriptomic deconvolution estimates and direct protein-level CD8 ADTs. In both modalities, GDF15^high^-CD8^low^ bins, defined as high GDF15 surrounded by low local CD8, were significantly more abundant in adenoma regions than in normal regions (post-hoc paired Wilcoxon, adj. *P* < 0.001 for both). Conversely, the GDF15^low^-CD8^high^ bins were significantly decreased in adenoma compared to normal regions (Figure 4H). Complementing this pattern-level analysis, we directly quantified local CD8+ T cell density and found that CD8 density measured by ADT was significantly lower around GDF15-high spots than GDF15-low spots within adenoma regions (paired Wilcoxon *P* = 0.018, Supplementary Figure 7H), but not in the normal (*P* = 0.15; Figure 4I). CD8 density around GDF15-high spots was also significantly decreased in advanced adenomas (paired Wilcoxon *P* < 0.001), but not in non-advanced adenomas (*P* = 0.31; Figure 4J). Together, these spatial analyses demonstrate that GDF15 expression in dysplastic epithelium is associated with local CD8+ T cell exclusion in an adenoma-specific and advancement-dependent manner.

### Single-cell spatial profiling links GDF15 to stemness and immune exclusion across the adenoma-to-carcinoma sequence

To spatially map the cellular and molecular landscape underlying colorectal tumorigenesis at the single cell level and validate the findings from the Visium CytAssist analysis, we constructed two tissue microarrays (TMAs) comprised of 111 cores from 16 patients who each had adenomas with co-occurring focal high grade dysplasia, intramucosal carcinoma or adjacent adenocarcinoma arising out of the adenoma, allowing analysis of matched normal colon mucosa, adenoma and adenocarcinoma (Figure 1, Supplementary Table 2). The TMA slides were subjected to spatial transcriptomic profiling using the Xenium Prime 5K platform, supplemented with 50 custom selected genes, with particular emphasis on genes associated with senescence and stemness. After quality control, 101 cores were used in downstream analysis. Prior to further analysis, PCA was conducted to assess for variation due to tissue type, colon location, gender, patient, Xenium slide, and age (Supplementary Figure 8).

Projection of tissue identity onto the UMAP revealed distinct yet partially overlapping distributions of cells derived from normal, adenoma, and cancer tissues (Figure 5A). Unsupervised clustering identified major cell types, assigned based on canonical marker gene expression (Figure 5B). Notably, normal samples included both tumor-adjacent tissues collected at surgical margins (designated “Adjacent N”). and independent normal tissue blocks (designated “Distant N”). GDF15 expression is highly enriched within epithelial cell clusters and low in stromal and immune compartments (Figure. 5C, Supplemental Figure 9A-B) and increases from normal to adenoma to cancer epithelium (Supplemental Figure 9C). These results indicate that epithelial cells are the predominant source of GDF15 early during tumorigenesis and that a subset of cancer cells also express high levels of GDF15. A closer annotation of the epithelial compartment identified 11 clusters with distinct features (C0–C10; Figure 5D and 5G). We found the relative abundance of many clusters shifted significantly across tissue types (Figure 5E, Supplemental Figures 11C-D). Normal tissues were predominantly composed of mature colonocytes (C4 and C5) and C2 goblet cells, with a smaller fraction of stem-like cells (C1 LGR5^+^ and C8 OLFM4□). In adenomas, we observed a marked expansion of C0 CCL5+/RNF43+ cells, C8 OLFM4□ cells and C10 CXCL14^+^ cells, accompanied by the emergence of C9 GDF15+ epithelial cells. In cancer, C3 CD55+ stress/EMT like cells and C6 proliferating cells expanded alongside decreases of C0 CCL5+/RNF43+ cells, C2 differentiated goblet cells, and C9 GDF15+ cells. Together, these shifts describe changes in epithelial states from normal through the adenoma-to-carcinoma sequence.

**Figure 5.**
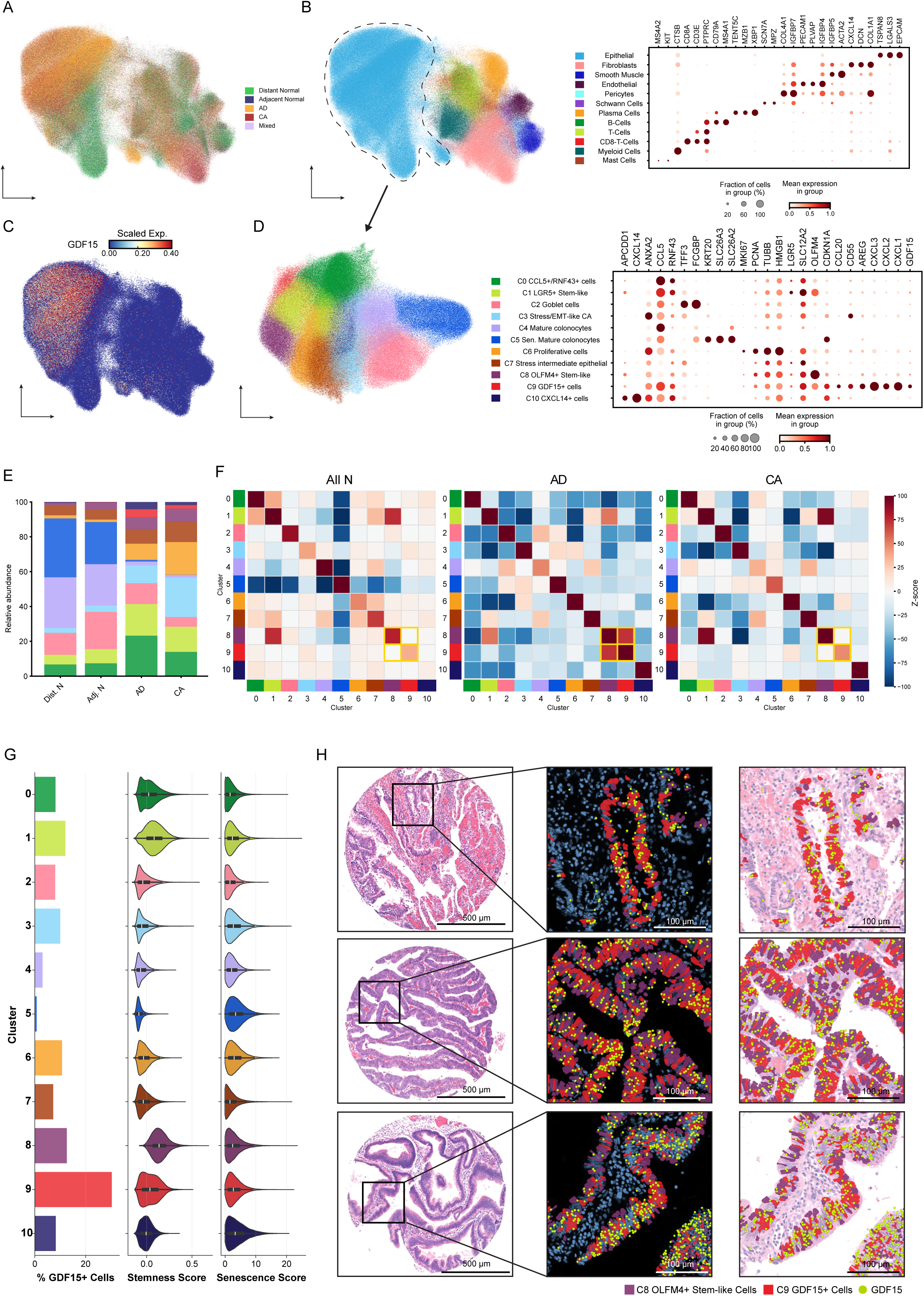
Spatially resolved characterization of colonic epithelial cells across normal, adenoma, and cancer TMA cores using Xenium 5K Prime reveals clusters with high GDF15 expression, SenePy scores, and stemness scores. (A) UMAP visualization of Louvain clustered cells (n= 797,861) from 101 Xenium cores (16 patients) colored by tissue type. (B) Left: UMAP visualization of all cells colored by cell-type annotation. Dotted line indicates epithelial cells. Right: Dotplot of representative marker genes used for annotation. Dot size represents the percentage of cells expressing each marker, and color represents the mean expression scaled per-gene. Full marker genes are shown in Supplemental Figure 10D. (C) UMAP visualization of all cells colored by normalized GDF15 expression. The color scale was clipped at the 1^st^ and 99^th^ percentiles to minimize the effect of outliers in visualization. (D) Left: UMAP visualization of subclustered epithelial cells (n=458,614) colored by annotated cluster. Right: Dotplot of representative marker genes used for epithelial cell type characterization. Dot size represents the percentage of cells expressing each marker, and color represents the mean expression scaled per-gene. Full marker genes are shown in Supplemental Fig. 11B. (E) Stacked barplot comparing the relative abundance of epithelial clusters across Distant Normal (78750 cells), Adjacent Normal (63441 cells), AD (208245 cells), and CA (108178 cells) tissue types. Per-patient compositional changes across tissue types are shown in Supplemental Figure 11A. (F) Heatmaps showing neighborhood enrichment of clustered epithelial cells for All Normal (combined Distant Normal and Adjacent Normal), AD, and CA tissue types. Color scale represents z-scores from neighborhood enrichment testing, and yellow boxes highlight the spatial relationship between C8 and C9 in each tissue type. (G) Barplot showing the percentage of cells within each epithelial cluster expressing at least 1 GDF15 transcript (Left). Violin plots showing the stemness score (Middle) and senescence scores (Right) for cells in each cluster. (H) H&E images of 3 representative AD cores with ROIs designated by black boxes (Left). OLFM4+ Stem-like epithelial cells (C8, purple), GDF15+ epithelial cells (C9, red), and GDF15 transcripts (green points) are visualized on DAPI stain (Middle) and H&E (Right) in ROIs using Xenium Explorer.

To assess spatial relationships among epithelial subtypes, we performed neighborhood analysis across epithelial subclusters (Figure 5F). In normal and cancer cores, the C1 LGR5+ stem-like cluster and C8 OLFM4+ stem-like cluster showed strong co-localization. Interestingly, a positive spatial association between C8 OLFM4□ stem-like cells and C9 GDF15□ cells appeared in adenoma. Their spatial organization within tissue architecture are demonstrated in representative adenoma cores (Figure 5H). These findings suggest the emergence of a spatially organized epithelial signaling niche centered on OLFM4□ and GDF15□ cells specifically in adenomas.

### Stemness and senescence programs co-localize within epithelial cell states

To further characterize functional programs within the epithelial compartment, we computed stemness and senescence scores based on established gene signatures and projected these onto the epithelial UMAP. Stemness scores were strongly enriched in cells belonging to C1 LGR5^+^ and C8 OLFM4□ stem/progenitor populations as well as C0 adenoma progenitor cells (Figure 6A, 6C, and 5D). Senescence scores were most strongly enriched in clusters C5, representing normal crypt-top colonocytes, and the stress/EMT associated clusters C3 and C10 (Figure 6B, 6C, and 5D). To further characterize epithelial stemness across tissue states, we examined the expression of each stemness marker used to calculate stemness score across the N-AD-CA sequence (Figure 6D). CDX2, a regulator of colon epithelial stemness^20^, was downregulated in both adenoma (AD) and carcinoma (CA) compared to normal tissue. OLFM4 showed the most pronounced increase in both expression level and cellular prevalence between distant normal and adjacent normal tissues, and this increase was sustained in AD and CA tissues, corroborating its central role in the stem-like expansion shown in Figure 5D–E. At the sample level, both stemness and senescence distributions showed gradual upward shifts, with increasing proportions of epithelial cells occupying higher-score bins across samples (Figure 6E–F). These patterns indicate continuous, heterogeneous activation of both programs across patients and tissue states.

**Figure 6.**
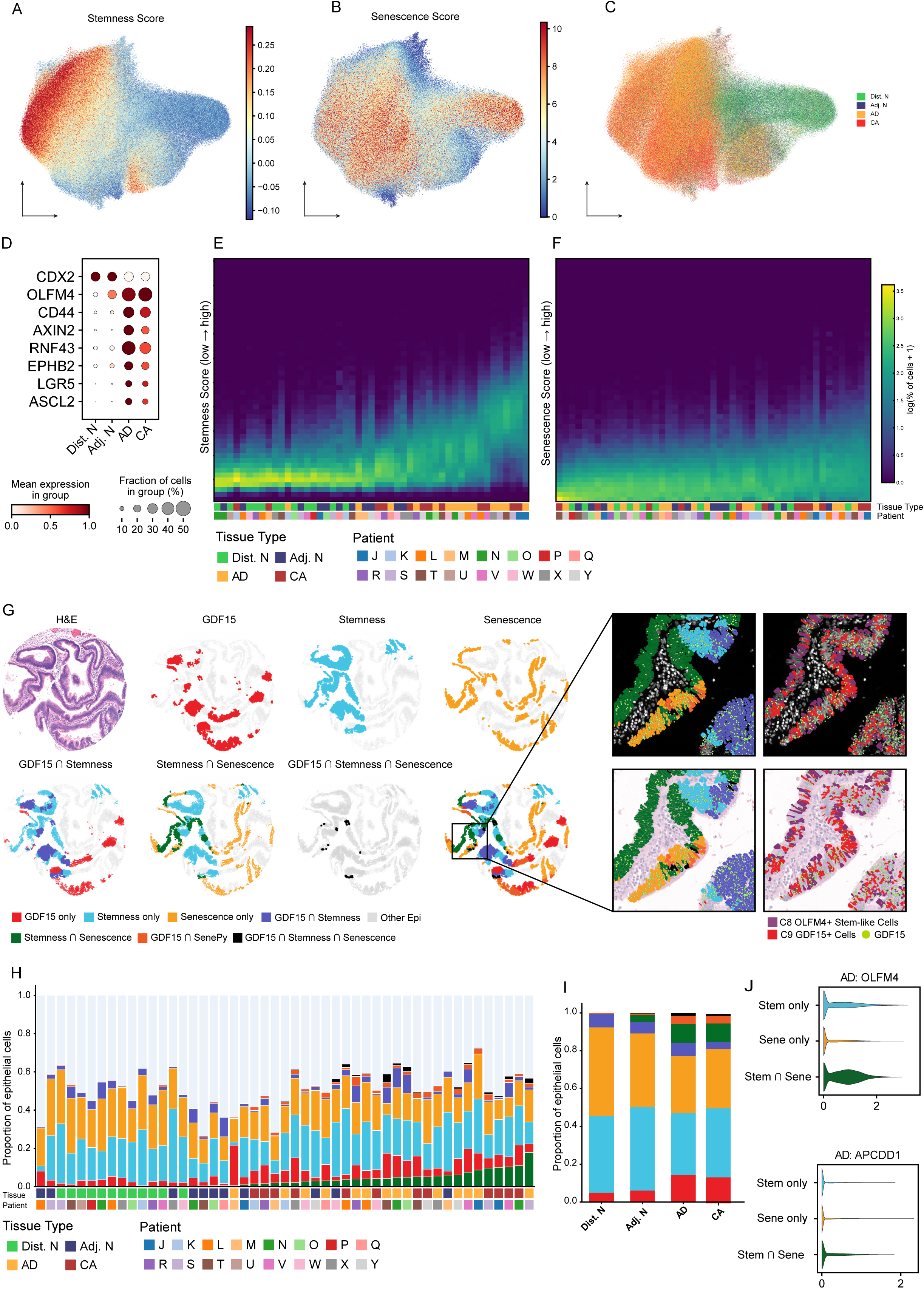
Stemness, senescence, and GDF15 features colocalize in a subset of adenoma and cancer cores. (A) UMAP visualization of Louvain clustered epithelial cells colored by stemness score. The color scale was clipped at the 1^st^ and 99^th^ percentiles to minimize the effect of outliers in visualization. (B) UMAP visualization of Louvain clustered epithelial cells colored by senescence score. The color scale was clipped at the 1^st^ and 99^th^ percentiles to minimize the effect of outliers in visualization. (C) UMAP visualization of Louvain clustered epithelial cells colored by tissue type. (D) Dotplot comparing expression of marker genes used to calculate the stemness score for stemness-high epithelial cells across different tissue types. Dot size represents the percentage of cells expressing each marker, and color represents the mean expression scaled per-gene. (E) Distribution of stemness score displayed as a heatmap where columns represent samples (all cells of a given tissue type for each patient) and rows represent 50 equally-sized bins of stemness score. The color scale represents the log1p-transformed percentage of epithelial cells falling within each bin. Samples are ordered by median score. Patient and tissue type labels are designated by color bars on the x axis. (F) Distribution of senescence score displayed as a heatmap where columns represent samples (all cells of a given tissue type for each patient) and rows represent senescence scores across all cells grouped into 50 equally sized bins. The color scale represents the log1p-transformed percentage of epithelial cells falling within each bin. Samples are ordered by median score. Patient and tissue type labels are designated by color bars on the x axis. (G) Top: GDF15, stemness score, and senescence score hotspots defined by the Local Moran’s I algorithm displayed on a representative adenoma core alongside the matched H&E core image. Bottom: Unions of GDF15, stemness score, and senescence hotspots. (H) Stacked barplot displaying the composition of GDF15, stemness, and senescence categories (defined by Local Moran’s I) within each tissue type for each patient. Samples are ordered by increasing overlap of stemness score and senescence hotspots. (I) Stacked barplot showing the composition of GDF15, stemness, and senescence hotspot categories (defined by Local Moran’s I) within each tissue type. Significance of hotspot changes between categories is detailed in Supplemental Figures 12B-C. (J) Violin plots comparing expression of transcripts OLFM4 (top) and APCDD1 (bottom) between Stemness only, Senescence Only, and Stemness ∩ Senescence hotspot cells in AD tissues. Violin plots comparing OLFM4 and APCDD1 expression across all hotspot groups are shown in Supplemental Figure 12E.

To spatially resolve the relationship between GDF15, stemness, and senescence at single-cell resolution, we applied Local Moran’s I (LISA) spatial statistic to define epithelial hotspots for each feature and mapped their pairwise and triple intersections across all cores (Figure 6G). Across patients and tissue types, the per-sample composition of these LISA categories varied substantially (Figure 6H), yet significant compositional changes between tissue types emerged (Figure 6I, Supplemental Figure 12B-C) In distant normal and adjacent normal epithelium, senescence hotspots were localized primarily in crypt-top colonocyte niches, reflecting the high senescence score of C5, whereas stemness hotspots occupied a spatially opposite niche likely representing the crypt bottom (Supplemental Figure 12A). There were almost no Stemness ∩ Senescence hotspots in either normal tissue type, and the few observed overlapping hotspots were found in adjacent normal tissue (Figure 6I). Interestingly, GDF15 ∩ Stemness hotspots were observed in both distant normal and adjacent normal tissue, while GDF15 ∩ Senescence hotspots were virtually nonexistent, suggesting stemness niches rather than senescence niches as the major source of GDF15 expression in normal epithelium.

In adenoma and cancer tissues, expansion of GDF15 hotspots was observed alongside increasing spatial overlap between GDF15, stemness, and senescence features. Stemness ∩ Senescence niches expanded most dramatically, marked by increased OLFM4 expression relative to Stemness-only and Senescence-only hotspots in AD and CA tissues (Supplemental Figure 12D and Figure 6J). In AD tissues, Stemness ∩ Senescence niches were also marked by a moderate increase in APCDD1 expression. Comparison of these two marker genes across all LISA categories revealed markedly increased OLFM4 expression in Stemness ∩ Senescence, GDF15 ∩ Stemness, and triple GDF15 ∩ Stemness ∩ Senescence niches in both AD and CA (Supplemental Figure 12E).

Together, these findings indicate that colorectal neoplasm progression is accompanied by the emergence of an epithelial niche integrating stemness and senescence-associated programs alongside an increase in GDF15-only and GDF15 ∩ Senescence hotspots.

### Spatial interaction between GDF15 and CD8A across normal, adenoma, and cancer tissues

Clustering of non-epithelial cells identified a T-cell population (Supplementary Figure 10), which was subclustered to identify a CD8+ cytotoxic cluster (Figure 7A and 7B) distinguished by selective co-expression of CD8A, CD8B, GZMA, GZMB, CCL4, and PRF1 relative to the other T cells (Figure 7C). To map the spatial relationship between GDF15 and CD8A, we aggregated Xenium transcripts into 30 μm² bins per core and applied bivariate Local Moran’s I (LISA), which identified CD8A^Low^-GDF15^High^ bins representing areas with low CD8A expression surrounded by areas with high GDF15 expression. Though some of these Low-High bins may represent epithelium surrounded by GDF15-high epithelium, visualization of Moran’s I categories alongside epithelial GDF15 expression and CD8-T-Cell location in each core revealed distinct regions where CD8-T-Cells are absent from the non-epithelial compartments immediately surrounding GDF15-expressing epithelium (Figure 7D). This pattern was most apparent in AD cores, where GDF15 expression was highest. Note that in CA cores, CD8 T cells were sparse throughout.

**Figure 7.**
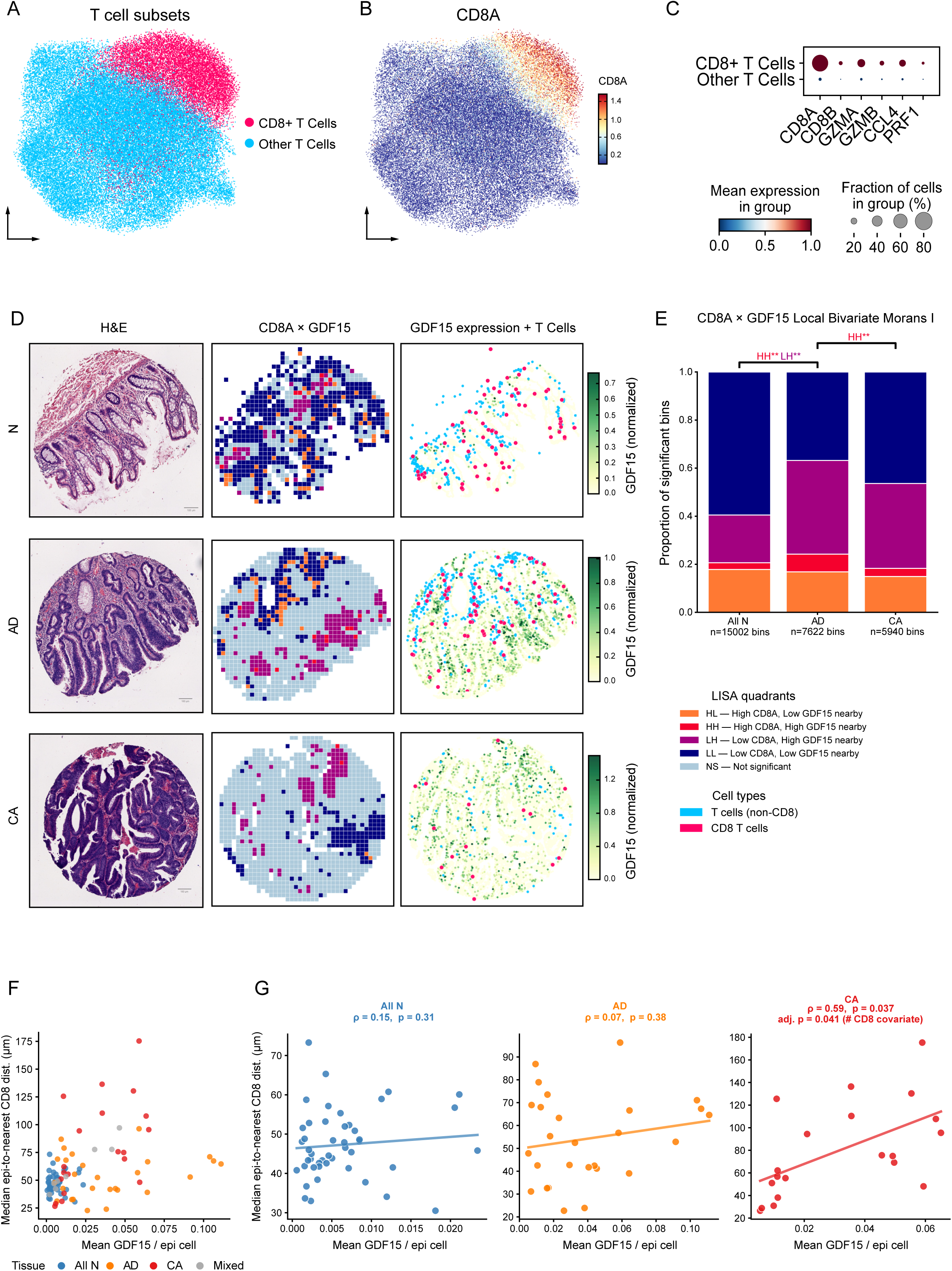
Spatial interaction between GDF15 and CD8A across normal, adenoma, and cancer TMA cores. (A) UMAP visualization of Louvain clustered T-cells. The CD8 T-cell annotated cluster is colored in pink, and all other T-cells are colored in blue. (B) UMAP visualization of Louvain clustered T-cells colored by CD8A expression. The color scale was clipped at the 1^st^ and 99^th^ percentiles to minimize the effect of outliers in visualization. (C) Dotplot of representative marker genes used to define the CD8 T-cell cluster. Dot size represents the percentage of cells expressing each marker, and color represents the mean expression scaled per-gene. (D) Left: H&E images of representative cores. Middle: Local bivariate correlation (LISA) of CD8A and GDF15 computed and displayed on 30um^2^ binned Xenium data. Significant quadrants (p < 0.05) were defined using 999 permutations. Right: Epithelial cells colored by normalized GDF15 expression alongside CD8 T-Cells (pink) and all other T-Cells (blue) (E) Stacked barplot displaying the composition of LISA quadrants across Normal (15,002 bins across 52 cores), AD (7622 bins across 35 cores) and CA (5940 bins across 22 cores) conditions. The Normal category represents combined Distant Normal and Adjacent Normal bins. Colors denote Moran’s I classes. **, P < 0.01. (F) Scatterplot displaying the median distance from each epithelial cell to the nearest CD8 T Cell vs. mean epithelial GDF15 expression for each core. Cores are colored by tissue type. Distant Normal and Adjacent Normal tissue types were combined into a single Normal tissue category, and cores with more than one epithelial tissue type were labeled as mixed. (G) Scatterplot displaying the median distance from each epithelial cell to the nearest CD8 T Cell vs. mean epithelial GDF15 expression for each core within each tissue type. Significance was determined by a rank-transformed linear mixed model using patient as the random intercept, though raw values (not ranks) and the OLS line are displayed for visualization purposes. BH-adjusted p-values and Spearman ρ values are reported for each tissue type. For CA cores, adj. P value represents the BH-adjusted p-value determined by a rank-transformed linear mixed model using patient as the random intercept and including # CD8 T Cells per core as a covariate.

Quantification across 15,002 normal, 7,622 adenoma, and 5,940 cancer bins confirmed these spatial patterns (Fig. 7E). Normal cores were dominated by Low-Low bins, reflecting globally low CD8A and GDF15 expression. In adenoma, both the High-High (high CD8A, high GDF15 nearby) and Low-High (low CD8A, high GDF15 nearby) categories were significantly expanded relative to both normal and cancer (post-hoc paired Wilcoxon, adj. P < 0.01 for both). Interestingly, we found the concurrent enrichment of High-High bins in adenoma, but not cancer.

To understand the core-level correlation of epithelial GDF15 expression and CD8 positioning, we compared the mean epithelial GDF15 expression for each core to the median distance from each epithelial cell to its nearest CD8-T-Cell across all cores (Figure 7F). Overall, normal cores exhibited the lowest mean epithelial GDF15 expression with lower median epithelial-to-CD8 distances, with mean GDF15 below 0.025 and median epithelial-to-CD8 distances generally below 60 μm, consistent with intact immune surveillance in non-neoplastic tissue. Adenoma cores spanned the widest range of GDF15 expression, showing moderate epithelial-to-CD8 distances. Cancer cores exhibited the greatest epithelial-to-CD8 distances across all tissue types, with several cores exceeding 120 μm. The relationship between epithelial GDF15 expression and epithelial-to-CD8 distance was tissue-context dependent (Figure 7G). In normal and adenoma cores, no significant association was detected (p = 0.31 and p = 0.38, respectively). In cancer, however, cores with higher mean epithelial GDF15 expression were significantly associated with greater epithelial-to-CD8 distances (adj. p = 0.037). This relationship in cancer cores remained significant after fitting a linear mixed model with per-core CD8 count as a covariate (adj. p = 0.041), indicating that CD8 exclusion from epithelial regions in cores with higher epithelial GDF15 expression is not merely a result of decreased CD8 density at the core level.

Altogether, these results suggest that GDF15-associated CD8 exclusion operates at two distinct spatial scales: a local, bin-level exclusion detectable in adenoma, and a broader, tissue-wide exclusion that gradually becomes significant in cancer.

## Discussion

A better understanding of the molecular and cellular changes that occur during colorectal cancer initiation and progression is critical for identifying targets for cancer prevention and early interception^21,22^. To date, studies have suggested certain cancer related hallmark behaviors play a role in the malignant transition of colorectal adenomas to cancer, including the development of neoangiogenesis (i.e. angiogenic switch)^23^, adoption of glutamine metabolism, and increased genomic instability due to TP53 loss, but a more complete analysis of the biological factors that associate with the malignant transformation of adenomas is lacking.

In this study, we analyzed two independent sample sets using complementary approaches: Visium CytAssist RNA and Protein co-detection across 22 nonadvanced and advanced tubular adenomas and Xenium Prime 5K across 101 patient-matched normal, adenoma, and carcinoma cores from 16 patients with colorectal adenomas with focal high-grade dysplasia or intramucosal carcinoma or adjacent invasive adenocarcinoma and also used public scRNA-seq data to validate our findings. Our studies provide evidence that senescence and stemness programs co-exist within a spatially defined epithelial niche and that expansion of that niche associates with colorectal adenoma progression. We find that GDF15, a prominent SASP factor^24^ may mediate the spatial coupling between these programs and that GDF15-high dysplastic epithelium is associated with local CD8+ T cell exclusion. These data suggest that, instead of being a static barrier to malignant transformation; cellular senescence in the epithelial cells paradoxically may also function as a spatially organized program that coordinates epithelial plasticity and immune evasion during adenoma progression.

Our finding that senescence co-occurs with stemness within the same epithelial niche is consistent with prior reports that the SASP can induce plasticity and stemness in neighboring cells^14,25,26^, a property increasingly recognized as a hallmark of cancer. We found that statistically controlling for GDF15 expression reduced the senescence–stemness spatial association in advanced adenomas but not in nonadvanced ones, which suggests that GDF15 is one mechanism by which these programs become coupled as lesions advance.

Our results add to a growing body of work on GDF15, a stress-induced cytokine and prominent component of the SASP^27,28^. Previous studies have demonstrated that tumor-derived GDF15 can block LFA-1-mediated T cell adhesion to endothelial cells, preventing T cell entry into the tumor microenvironment, and serum GDF15 levels predict anti-PD-1 failure in cancer patients^28,29^. In colorectal cancer, GDF15 has been shown to originate from senescent peritumoral epithelial cells in normal-adjacent tissue, where it sustains tumor growth through a metabolic feedback loop^30^. Interestingly in the normal tissue adjacent to the lesion, the expression patten of GDF15 was found to be correlated with the OLFM4 expression. In adenoma and cancer tissue, GDF15 expression was upregulated, and CD8+ T cell density was lower around GDF15-high spots than GDF15-low spots but not in normal tissue. This exclusion effect was strongest in advanced adenomas and not significant in nonadvanced adenomas. In cancer cores, GDF15 expression correlated with epithelial-to-CD8 distance across the full cohort, suggesting that the areas with GDF15-related CD8 exclusion may have expanded during the adenoma to cancer sequence. GDF15 signals through GFRAL and blocking this axis has been shown to restore T cell trafficking in clinical studies^29^. Our results raise the possibility that a similar approach could restore CD8+ T cell access to the dysplastic niche in the adenoma setting, although this remains to be tested directly. Our data are correlational, and functional studies are needed before causality can be established.

Our study has several limitations. First, Visium spots contain multiple cells, and although we used spatial deconvolution, senescence and stemness programs cannot be assigned to individual cell types with full confidence. Second, the Xenium TMA format samples discrete cores rather than whole tissue sections, so spatial context beyond each core is not captured. Third, our correlation analysis supports GDF15 as a mediator of senescence–stemness coupling but cannot establish causality on its own. Functional validation in vitro or in vivo models is needed to establish causality. Finally, our cohorts are cross-sectional; longitudinal sampling of the same lesion over time would be invaluable to confirm our findings.

Overall, our studies provide evidence that senescent epithelial cells form a spatially organized signaling niche during colorectal adenoma progression, and that GDF15, as a central SASP factor in this niche, can mediate the senescence-stemness-immune evasion axis during adenoma to cancer progression. This niche is present in early lesions and expands as adenomas advance toward carcinoma. Our findings warrant further investigation of GDF15 as a potential target for colorectal cancer prevention and interception.

## Methods and Materials

### Study cohort and pathological annotation

Twenty-four FFPE colon tubular and tubulovillous adenoma specimens were included, stratified by lesion size and histology into nonadvanced adenomas (aka small; diameter <10 mm, n = 8) and advanced adenomas^31^ (aka large; diameter >10mm, n = 16) (Supplementary Table 1). All cases were reviewed and annotated by two independent pathologists. For spatial transcriptomics, high-resolution H&E images corresponding to Visium capture areas were reviewed by a board-certified pathologist using Loupe Browser (10x Genomics, v9.0.0). Each spot was manually classified as stroma, normal epithelium, or low-grade dysplasia based on histomorphology, with the pathologist blinded to RNA expression, antibody-derived tag (ADT) expression, and computational clustering results.

Of note, all tissue samples were de-identified. The studies were subjected to Fred Hutchinson Cancer Center IRB review and were approved.

### Tissue microarray construction and pathological annotation

Two tissue microarray (TMA) blocks were constructed comprising 111 patient-derived cores from 16 patients, representing normal colon, tubular adenoma, and colorectal cancer tissues. Target tissues were sampled in replicate using 1-mm diameter cores. Tonsil and normal colon reference tissues were included on each TMA block as standardized controls to perform normalization and batch correction across spatial transcriptomics runs. The Xenium Prime 5K panel was supplemented with 50 custom genes selected to capture epithelial, stromal, and immune compartments, with emphasis on senescence, inflammation, and epithelial plasticity pathways. Following Xenium data acquisition, cores were reviewed and annotated by a board-certified pathologist. Cores with fewer than 30 epithelial cells after epithelial masking were excluded from spatial analyses.

### Immunohistochemistry staining

Immunohistochemistry (IHC) was performed on consecutive sections from Visium-profiled adenoma specimens using a Ventana Discovery Ultra Autostainer IHC (Roche Diagnostics, Basel, Switzerland) for p21 Waf1/Cip1 (12D1) (Cell Signaling, Cat. No. 2947), p16 INK4a (Biocare Medical, Cat. No ACI3231A), and Ki-67 (D2H10) (Cell Signaling, Cat. 9027). IHC was performed using the DISCOVERY anti-HQ HRP kit. The slides were steamed for 48 minutes in Cell Conditioning 1 (CC1) solution (Roche Diagnostics, Cat. No. 950-500) for antigen retrieval. Then the primary antibody was applied for 1 hour at p21 (1:50), p16 (1:50), and Ki-67 (1:400).

### Spatial library generation and sequencing

Spatial gene expression libraries were generated using 10x Genomics Visium Spatial Gene Expression (RNA only; n = 4 slides) and Visium CytAssist Spatial Gene and Protein Expression (RNA + ADT; n = 20 slides), with 31 antibodies and 4 isotype controls captured as ADTs. Libraries were sequenced on an Illumina platform and raw sequencing data processed with Space Ranger (v3.0) aligned to the GRCh38 reference genome. Two samples failing Space Ranger quality control were excluded from downstream analyses.

### Quality control

Spot-level RNA quality control was applied in Seurat (v5.3.0) in R (v4.4.0). Spots were excluded if they met any of the following criteria: mitochondrial RNA fraction ≥40%, total UMI count ≤300, or location outside the tissue area as determined by the Space Ranger/Loupe tissue mask. For ADT data, spots were removed if the isotype control proportion was ≥20% or fewer than 20 antibodies were detected; these spots were retained as RNA-only for subsequent RNA analyses.

### Normalization, dimensionality reduction, and multimodal integration

RNA counts were normalized within each spot for library size, log-transformed, and scaled. We identified the 2,000 most highly variable genes, which were used for principal-component analysis and downstream FastMNN integration. ADT counts were normalized using dsb (v2.0.0) with isotype controls as background, performed within each slide, then scaled and batch-corrected across samples using Harmony (v1.2.4). RNA and ADT modalities were integrated using Seurat’s weighted nearest-neighbour (WNN) framework, incorporating the first 30 RNA PCs and first 15 ADT PCs, to learn per-spot modality contributions. The resulting WNN graph was used for UMAP visualization and Louvain clustering using the smart local moving algorithm.

For RNA-only spots originating from RNA+ADT slides (after ADT quality control filtering), transfer anchors were learned from the integrated RNA+ADT object to map these spots into the same low-dimensional embedding and transfer cluster labels. RNA-only library samples were processed with the same RNA pipeline and mapped to the integrated reference using Seurat anchor-based transfer learning to ensure cross-dataset comparability.

### Cluster annotation

Cluster marker genes were identified using Seurat FindAllMarkers(). For each sample–condition stratum, cluster counts were converted to relative abundances, with absent clusters assigned a proportion of zero. Overall compositional differences were assessed by PERMANOVA with 999 permutations on Hellinger-transformed proportion matrices using vegan (v2.7-2), and homogeneity of multivariate dispersion was evaluated using betadisper() and permutation tests. Cluster-specific differences were assessed using Wilcoxon tests with Benjamini–Hochberg false-discovery-rate correction. Comparisons between pathologist-annotated Normal and Adenoma regions were restricted to samples containing both regions, with cluster-specific tests paired within sample. Comparisons between nonadvanced and advanced adenomas were performed as unpaired sample-level tests. Clusters were annotated by integrating RNA marker expression, ADT expression, and spatial localization relative to pathologist-annotated tissue regions.

### Spatial pseudotime and diffusion map analyses

Spatial pseudotime was inferred using stLearn (v1.2.2) in Python. Analyses were restricted to a high-confidence epithelial compartment defined by the intersection of epithelial-like clusters and pathologist annotations (normal epithelium and adenoma), excluding stromal regions. The pseudotime root was anchored to the crypt-base epithelial cluster (cluster 1) by selecting the spot closest to the median UMAP coordinate of that cluster. Spatial localization prior to pseudotime inference was performed using DBSCAN-based spatial clustering, with slide-adaptive epsilon estimated from spot-to-spot array distances. For slides containing disconnected tissue islands, spots were partitioned by connected components of a k-nearest neighbour (kNN) graph (k = 8) and pseudotime computed independently per island. Pseudotime values were scaled from 0 to 1 within each sample or island.

To visualize the continuous latent structure of the epithelial compartment, diffusion map dimensionality reduction was performed using the destiny R package (v3.20.0) on the MNN-reduced embedding with a fixed Gaussian kernel width (sigma = 1). For two-dimensional visualization, diffusion components 1 and 3 were displayed because this projection more clearly resolved the branching epithelial-state structure.

### Cell cycle scoring and developmental potential estimation

Spot-level cell cycle status was estimated using two complementary methods: Seurat CellCycleScoring() with canonical S-phase and G2/M gene sets, and ccAFv2 (v0.0.0.9) in spatial mode (probability threshold = 0.5). Absolute developmental potential was estimated using CytoTRACE 2 (v1.1.0) with default parameters; scores range from 0 (terminally differentiated) to 1 (totipotent).

### Gene set enrichment analysis

Pathway enrichment was assessed at the cluster level using four algorithms, including AUCell, UCell, singscore, and JASMINE, implemented in irGSEA (v3.3.3). Results were integrated across methods using Robust Rank Aggregation (RRA); gene sets with RRA p < 0.05 were considered robustly enriched. MSigDB Hallmark and SenMayo gene sets were obtained from MSigDB; Immune exclusion signature (IEX) and stemness signatures were based on published gene sets and used in their original form (Supplementary Table 3)^4,12^. UCell scores are reported for visualization.

### Senescence scoring

Spot-level senescence scores were computed using SenePy (v1.0.1) score_hub() with the universal hub module. Senescent outlier spots (SenePy^high^) were defined as those with scores exceeding two standard deviations above the mean, following the original SenePy publication.

### Luminal surface distance analysis

Luminal surface spots were manually annotated on H&E-aligned Visium sections. Disconnected tissue islands were separated using semla (v1.4.0), and distances from each non-stromal spot to the nearest annotated luminal surface spot were calculated within each island. Pixel distances were converted to micrometers using the median nearest-neighbor Visium spot spacing (100-µm center-to-center). Annotated surface spots served only as spatial anchors and were excluded from trend fitting. To visualize changes in stemness and senescence scores along the crypt axis, spots were grouped into 100-µm bins according to their distance from the luminal surface. For each sample, tissue region, and distance bin, we calculated the mean score and number of contributing spots. Bin-level mean scores were visualized using generalized additive model smoothing with a cubic regression spline, fitted separately for normal and adenoma regions. Shaded bands represent the 95% confidence intervals of the fitted mean trends.

### Senescence–Stemness–Immune Exclusion Co-occurrence Score

To quantify the spatial co-occurrence of senescence, stemness, and immune exclusion, we developed a Senescence–Stemness–Immune Exclusion co-occurrence score (SSIE co-occurrence score). Because the three component scores were derived using different methods and had different numerical scales, each score was independently converted to a within-sample percentile rank. For each spatial location, percentile ranks were calculated as ((r-0.5)/n), where (r) is the average rank of the score within the sample and (n) is the number of spatial locations with a non-missing value. Higher percentile values therefore represented stronger relative enrichment of the corresponding state within each sample.

The SSIE co-occurrence score was defined as the geometric mean of the percentile-ranked senescence, stemness, and immune-exclusion scores:

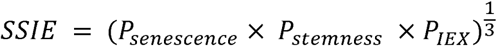

The geometric mean penalizes low values in any individual component and therefore produces a high score only when senescence, stemness, and immune exclusion are concurrently elevated. Because percentile ranking was performed separately within each sample, the resulting score represents relative spatial co-enrichment within each specimen rather than an absolute measure for direct comparison of score magnitude across samples.

### Differential expression analysis

Exploratory spot-level differential expression analyses were performed on spatially defined epithelial regions. Three comparisons were conducted: (i) advanced versus nonadvanced adenoma spots within dysplastic clusters C2 and C4; (ii) stemness-high versus stemness-low spots within the same dysplastic clusters; and (iii) senescence-high spots from dysplastic epithelial regions versus senescence-high spots from normal epithelial regions. Genes with adjusted P < 0.05, log fold-change > 0.2, and a difference in detection frequency > 10% were considered enriched in each comparison. Genes enriched across all three contrasts were intersected and filtered for genes encoding secreted proteins.

For the stemness comparison, stemness-high and stemness-low spots were defined as those with scores above the pooled 75th percentile and below the pooled 25th percentile, respectively; intermediate spots were excluded. Senescence-high spots were defined as senescent outlier spots using the SenePy criterion described above. For GDF15, expression was additionally summarized at the sample level. Advanced and nonadvanced adenomas were compared using a two-sided Wilcoxon rank-sum test, whereas stemness-high versus stemness-low dysplastic regions and senescence-high dysplastic versus senescence-high normal epithelial regions were compared using two-sided paired Wilcoxon signed-rank tests.

### Local Moran’s I analysis

To evaluate pairwise spatial co-associations between spot-level gene expression, ADT levels, and pathway scores, bivariate local Moran’s I (LISA) was computed using rgeoda (v0.1.0). For each Visium section, spatial weights were constructed using a kNN graph (k = 6) based on spot coordinates. Statistical significance was assessed by permutation (999 permutations; two-sided; p < 0.05). Significant spots were classified as High–High, Low–Low, High–Low, or Low–High, with non-significant spots labelled "Not significant", and were excluded from quadrant composition calculations and subsequent statistical comparisons.

For bivariate analysis of CD8A and GDF15 in Xenium data, cells were aggregated into 30 µm^2^ bins by floor division, with total GDF15 and CD8A transcript counts computed per bin. Bin tissue type labels were assigned as the majority epithelial tissue type within each bin; non-epithelial bins in mixed cores were assigned the label of their nearest epithelial-containing bin. Bivariate local Moran’s I was applied per core with CD8A as the focal feature and GDF15 as the lag feature (999 permutations, p < 0.05). This yielded four categories of significant bins: High–High (high CD8A, high neighboring GDF15), High–Low, Low–Low, and Low–High (low CD8A, high neighboring GDF15). Compositional differences of LISA classes across tissue types were assessed by PERMANOVA (vegan v2.7.2, 999 permutations), with dispersion homogeneity verified by betadisper(). Category-level post-hoc tests used Wilcoxon rank-sum or paired signed-rank tests with BH correction.

For Xenium data, Moran’s I was additionally computed per core using a 50-µm neighborhood radius and 999 permutations (esda v2.7.0, PySAL)^32^. Significant (p < 0.05) High–High and Low–High cells were classified as hotspots for each feature (GDF15, stemness score, and senescence score). Hotspot unions were computed across features within each core.

### Spatial deconvolution

Cell type abundances within Visium spots were estimated using Cell2location (v0.1.5) against a curated single-cell colon adenoma reference atlas, restricted to colonoscopy-derived tubular and tubulovillous adenoma samples. Neoplastic epithelial populations in the reference were relabeled with a "Dysplastic" prefix to accurately reflect the precancerous nature of the cohort and distinguish them from carcinoma-derived cells. Raw count matrices were supplied for both the Visium data and the single-cell reference. Mitochondrial, ribosomal, and hemoglobin genes were removed from the reference, Cell2location gene filtering was applied (cell count cutoff = 5, cell percentage cutoff = 0.03, nonzero mean cutoff = 1.12), and the reference and Visium matrices were restricted to their intersecting genes. The reference regression model was trained for 500 epochs, and the spatial mapping model was trained for 5,000 epochs with a batch size of 2,500, using a prior of 10 cells per spot and detection alpha = 20. Posterior abundance was estimated using 1,000 samples, and the 5th percentile of the posterior distribution was used for downstream spatial analyses.

### Partial spatial correlation analysis

To evaluate whether GDF15 mediates the spatial co-association between senescence and stemness programs, we computed spatial lag correlations within each Visium adenoma sample. For every spot, a spatial lag of stemness score was defined as the average stemness score of its nearest neighboring spots (k = 6). We then calculated Spearman correlations between spot-level senescence scores and the corresponding lagged stemness scores, yielding one spatial lag correlation coefficient per sample.

To obtain partial spatial correlations, spot-level senescence and stemness scores were each adjusted for GDF15 expression within each adenoma sample. We then recomputed the spatial lag correlation between GDF15-adjusted senescence at each spot and GDF15-adjusted stemness averaged across its six nearest neighboring spots. Correlation coefficients were Fisher Z-transformed, and changes in spatial association before versus after GDF15 adjustment were evaluated using paired Wilcoxon signed-rank tests and a sign-flip permutation test (999 permutations), overall and stratified by advanced versus non-advanced adenoma.

### CD8+ T cell spatial neighborhood analysis

GDF15 expression was quantified from Visium RNA, and CD8+ T cells were identified using normalized ADTs. Within each sample, CD8-positive epithelial spots were defined using a composite T-cell gate (CD3E, CD8A, CD45RA, and CD45RO) to minimize false positives. Analyses were restricted to epithelial-enriched clusters, excluding stromal spots, and performed within individual tissue islands to avoid cross-island artefacts. For each epithelial source spot, local CD8 density was computed as the fraction of nearby epithelial spots (within a fixed two-spot-spacing circular radius) that were CD8-positive, excluding the center spot. Two comparisons were performed: (i) CD8 density around GDF15-high spots in adenoma versus normal epithelium; and (ii) CD8 density around GDF15-high versus GDF15-low spots within low-grade dysplastic and normal epithelial regions separately, using within-region median binning to control for regional expression differences. All tests were performed at the sample level using paired Wilcoxon signed-rank tests.

### Single-cell RNA-seq analyses

Public single-cell RNA-seq data from human colorectal adenomas were obtained from a prior publication^16^. Datasets A and B correspond to the two adenoma scRNA-seq cohorts (“discovery” and “validation”) reported by Chen et al^4^. Both are analyzed to demonstrate that the senescence and stemness programs map to distinct dysplastic epithelial subpopulations across independent cohorts. Original dimensionality reduction and cell type annotations were retained; only tubular adenoma samples were included. For analyses of epithelial subpopulations, data were re-normalized, scaled using HVFs, and reclustered in Seurat to define epithelial subclusters.

### Xenium Data preprocessing

Xenium TMA data from two slides were read into Python as SpatialData^33^ (v0.4.0) objects using spatialdata-io (v0.2.0) and stored in Zarr format. Cells with fewer than 15 transcripts or 3 detected genes were excluded, as were genes expressed in fewer than 3 cells. Cells below the 1st percentile of average DAPI intensity (from the Xenium morphology image) were additionally removed. Per-cell counts were normalized to a target of 100 and log1p-transformed. PCA was computed at the core level using scikit-learn^34^ (v1.7.1) with 10 components.

### Xenium Cell clustering and annotation

The two Xenium TMA datasets were concatenated into a single AnnData^35^ (v0.11.4) object and clustered using Scanpy’s^36^ Louvain algorithm (16 nearest neighbours, 50 PCs, resolution = 0.6). Batch effects attributable to TMA slide and patient were corrected using Harmony^37^ (harmonypy v0.0.10, theta = 0.5 for each effect). This yielded 11 clusters, of which six were identified as epithelial based on EPCAM expression. A single intermediate cluster with low transcript counts was excluded. Marker genes for each cluster were identified using Scanpy’s rank_genes_groups (Wilcoxon method) on the normalized layer and visualized as dot plots with per-gene scaled expression. Xenium Explorer (v4.1.0) was used to generate visualizations showing the colocalization of clusters with specific transcripts as overlays on both DAPI stain and H&E background. H&E images were aligned to the Xenium DAPI channel using the manual keypoint alignment feature.

### Epithelial clustering

Epithelial cells were split by tissue type (Distant Normal, Adjacent Normal, Adenoma, Cancer) and each subset independently subclustered (30 nearest neighbors, 50 PCs, resolution = 0.8) with Harmony batch correction. Fibroblasts initially misannotated as epithelial when clustering all cells were identified in the Distant Normal and Adjacent Normal subsets and removed. All remaining epithelial cells were jointly reclustered using 20 nearest neighbors and 50 PCs with Harmony correction. Final cell labels were corrected using the IDs of the identified misannotated fibroblasts. Compositional differences across tissue types were statistically validated using Mann-Whitney U tests for each cluster and tissue type comparison on cells aggregated to the patient and tissue type level. Due to the compositional nature of this data, scCODA (v0.1.9) was also used to identify credible compositional changes between tissue types. Cluster 7 was used as the reference cluster, and cells were again aggregated to the patient and tissue type level.

### Non-epithelial clustering

Non-epithelial cells were subset and reclustered (resolution = 0.2), yielding two stromal and two immune clusters. Immune cells were reclustered (resolution=0.6) and annotated using the following markers: T cells (CD3E, PTPRC); myeloid cells (CTSB, SLC40A1, LIPA, PTPRC); B cells (MS4A1, CD19); plasma cells (XBP1, MZB1, TENT5C); and mast cells (KIT, MS4A2). Stromal cells were reclustered (resolution=0.8) and annotated as fibroblasts (COL1A1, COL5A1, POSTN, DCN); endothelial cells (PECAM1, PLVAP); Schwann cells (MPZ, SCN7A, L1CAM); smooth muscle cells (ACTA2, MYLK, GREM2); and pericytes (COL4A1, COL4A2, MCAM).

### T-cell subclustering

T cells were subclustered using 30 nearest neighbors and 50 PCs, with Harmony batch correction applied for patient (theta = 0.5). A CD8+ cytotoxic T cell cluster was defined by expression of CD8A, CD8B, GZMA, GZMB, CCL4, and PRF1.

### Stemness and senescence scoring

Epithelial cells were scored for senescence using the intestine_epithelial_0 SenePy module. Stemness scores were computed using Scanpy’s score_genes function using the available genes from above-mentioned stemness signature: CD44, AXIN2, RNF43, EPHB2, CDX2, LGR5, OLFM4, and ASCL2. Score distributions were visualized as heatmaps, where cells were grouped into 50 equally sized bins and the log1p-transformed percentage of epithelial cells per bin plotted across samples ordered by median score.

### Spatial neighborhood enrichment analysis

Spatial neighborhood graphs were computed per core and per tissue type using Delaunay triangulation in Squidpy^38^ (v1.6.5). For each tissue type, core-level neighborhood enrichment matrices were concatenated and reordered to match the corresponding anndata objects. Neighborhood enrichment between epithelial clusters was computed using Squidpy (1,000 permutations); color scales represent Z-scores clipped at [−100, 100].

### Epithelial GDF15–CD8+ T cell distance analysis

For each core, mean epithelial GDF15 expression and the median distance from each epithelial cell to its nearest CD8+ T cell were calculated. Distant Normal and Adjacent Normal tissue types were pooled into a single Normal category; cores with mixed epithelial tissue types were labelled "mixed". For each tissue type, the relationship between mean epithelial GDF15 expression and epithelial-to-CD8 distance was evaluated using a rank-transformed linear mixed model with patient as a random intercept (statsmodels v0.14.5)^39^: rank(distance) ∼ rank(GDF15) + (1 | patient). Adjusted p-values were computed using BH correction. To confirm that significant results were not confounded by the number of CD8 T-Cells per core, the CA cores were also tested with CD8 count as a covariate and patient as a random intercept: rank(distance) ∼ rank(GDF15) + rank(CD8 count) + (1 | patient).

### GDF15 expression comparisons across cell and tissue types

GDF15 expression was compared between epithelial and non-epithelial cells, and between epithelial cells and fibroblasts within tissue types, using two-sided paired Wilcoxon signed-rank tests paired by patient (Scipy v1.15.2). GDF15 expression was additionally compared across epithelial tissue types using two-sided paired Wilcoxon tests. BH correction was applied for multiple comparisons (statsmodels v0.14.5). Cores with mixed epithelial tissue types were excluded from tissue-type comparisons.

### Statistical comparisons of continuous metrics

Continuous per-spot metrics were compared at the sample level to avoid treating individual spots as independent observations. Spot-level scores were first aggregated per sample (median). For spot-level conditions defined within a sample, per-sample aggregated scores were compared using paired Wilcoxon signed-rank tests. For sample-level conditions, groups were compared using unpaired Wilcoxon rank-sum tests. Effect sizes are reported as the median difference between conditions based on per-sample aggregated scores. For multiple comparisons, the Benjamini-Hochberg (BH) adjusted p-values were reported.

### Compositional analysis

Compositional differences in spot category proportions between conditions were assessed by summarizing each sample into category proportions, applying Hellinger transformation, and computing Euclidean distances. Overall compositional differences were tested by PERMANOVA (vegan v2.7.2, 999 permutations). Homogeneity of within-group dispersions was verified using betadisper() followed by permutation testing (999 permutations). Category-level post-hoc tests were performed using Wilcoxon rank-sum tests (sample-level conditions) or paired Wilcoxon signed-rank tests (within-sample conditions), with BH correction applied across categories.

### Statistical analysis

All statistical tests were two-sided unless otherwise specified, and P < 0.05 was considered statistically significant. The biological sample or patient, as appropriate to the study design, was treated as the independent unit for confirmatory inference. Where spot-, cell- or bin-level analyses were used, within-sample or within-patient dependence was addressed by sample-level aggregation, matched tests or mixed-effects models, as specified in the relevant subsections; exploratory spot-level differential-expression analyses are explicitly identified. Continuous spot-level outcomes were aggregated per sample and compared using paired Wilcoxon signed-rank tests for matched conditions or Wilcoxon rank-sum tests for independent groups. Compositional outcomes were assessed by PERMANOVA after Hellinger transformation, with homogeneity of dispersion evaluated by permutation testing; category-level post-hoc comparisons used the corresponding paired or unpaired Wilcoxon tests. Patient-level dependence in Xenium analyses was addressed by paired testing or mixed-effects models with patients as a random intercept, as specified in the relevant subsections. P values were adjusted for multiple comparisons using the Benjamini–Hochberg method where applicable. Spatial statistics, permutation counts, effect-size definitions and analysis-specific thresholds are described in the relevant subsections. Analyses were performed in R (v4.4.0) and Python (v3.10.18); key package versions are specified in the relevant subsections. Data visualization used ggplot2, matplotlib and seaborn.

## Ethics statement and approval

All samples included in this study are all de-identified and that IRB committee at Fred Hutchinson Cancer Center has approved the research.

## Data Availability

The Visium spatial transcriptomic data generated in this study have been deposited in the Gene Expression Omnibus under accession GSE342139. The Xenium in situ transcriptomic data have been deposited in GEO under accession GSE341852. These records are currently available to editors and reviewers through confidential reviewer-access tokens and will be released publicly upon publication. Source data are provided with this paper.

## Code Availability

Analysis code and pipelines are hosted at https://github.com/grady-lab/Spatial-multi-omics.

## Acknowledgements

We thank the patients who generously donated tissue samples for this study. We also thank the Experimental Histopathology, Genomics & Bioinformatics, and Spatial Genomics Shared Resources at Fred Hutchinson Cancer Center for their technical support.

## Author contributions

M.Y. and W.M.G. conceived and supervised the study. M.Y. and W.M.G. acquired funding. M.Y. oversaw the workflow, data generation, and quality control. Y.X. and L.A. performed formal analysis and generated figures for the Visium and Xenium datasets, respectively. K.C. contributed to sample acquisition and tissue processing. E.D. contributed to sample acquisition and the design of the TMA blocks and was responsible for constructing the TMA blocks. L.C. and C.K. contributed to bioinformatic and statistical analysis. K.A.M. performed pathological review and annotation of Visium and Xenium tissue sections. A.M.D. and J.H. oversaw/performed immunohistochemistry. W.M.G. and D.R. contributed to sample acquisition, and pathology review. E.W.N. contributed to spatial multiomic profiling and analysis of T cells; W.S. contributed to biostatistical methodology/analysis. M.Y., Y.X., L.A. wrote the original draft. All authors reviewed and edited the manuscript and approved the final version.

## References

1 Fearon, E. R. & Vogelstein, B. A genetic model for colorectal tumorigenesis. Cell 61, 759–767 (1990). 10.1016/0092-8674(90)90186-I

2 Muzny, D. M. et al. Comprehensive molecular characterization of human colon and rectal cancer. Nature 487, 330–337 (2012). 10.1038/nature11252

3 Becker, W. R. et al. Single-cell analyses define a continuum of cell state and composition changes in the malignant transformation of polyps to colorectal cancer. Nature Genetics 54, 985–995 (2022). 10.1038/s41588-022-01088-x

4 Chen, B. et al. Differential pre-malignant programs and microenvironment chart distinct paths to malignancy in human colorectal polyps. Cell 184, 6262–6280.e6226 (2021). 10.1016/j.cell.2021.11.031

5 Lu, Z. et al. Polyclonal-to-monoclonal transition in colorectal precancerous evolution. Nature 636, 233–240 (2024). 10.1038/s41586-024-08133-1

6 Campisi, J. & d’Adda di Fagagna, F. Cellular senescence: when bad things happen to good cells. Nature Reviews. Molecular Cell Biology 8, 729–740 (2007). 10.1038/nrm2233

7 López-Otín, C., Blasco, M. A., Partridge, L., Serrano, M. & Kroemer, G. The hallmarks of aging. Cell 153, 1194–1217 (2013). 10.1016/j.cell.2013.05.039

8 Hoi, X. P. et al. Cellular senescence in precancer lesions and early-stage cancers. Cancer Cell 44, 6–11 (2026). 10.1016/j.ccell.2025.10.006

9 Park, S. S. et al. Cellular senescence is associated with the spatial evolution toward a higher metastatic phenotype in colorectal cancer. Cell Reports 43, 113912 (2024). 10.1016/j.celrep.2024.113912

10 Sun, J. et al. A Glb1-2A-mCherry reporter monitors systemic aging and predicts lifespan in middle-aged mice. Nature Communications 13, 7028 (2022). 10.1038/s41467-022-34801-9

11 Colucci, M. et al. Senescence in cancer. Cancer Cell 43, 1204–1226 (2025). 10.1016/j.ccell.2025.05.015

12 Heiser, C. N. et al. Molecular cartography uncovers evolutionary and microenvironmental dynamics in sporadic colorectal tumors. Cell 186, 5620–5637.e5616 (2023). 10.1016/j.cell.2023.11.006

13 Sanborn, M. A., Wang, X., Gao, S., Dai, Y. & Rehman, J. Unveiling the cell-type-specific landscape of cellular senescence through single-cell transcriptomics using SenePy. Nature Communications 16, 1884 (2025). 10.1038/s41467-025-57047-7

14 Milanovic, M. et al. Senescence-associated reprogramming promotes cancer stemness. Nature 553, 96–100 (2018). 10.1038/nature25167

15 Milanovic, M., Yu, Y. & Schmitt, C. A. The Senescence-Stemness Alliance - A Cancer-Hijacked Regeneration Principle. Trends in Cell Biology 28, 1049–1061 (2018). 10.1016/j.tcb.2018.09.001

16 Marteau, V. et al. Single-cell integration and multi-modal profiling reveals phenotypes and spatial organization of neutrophils in colorectal cancer. Cancer Cell 44, 146–165.e114 (2026). 10.1016/j.ccell.2025.12.003

17 Li, S. et al. Advancing biological understanding of cellular senescence with computational multiomics. Nature Genetics, 1–14 (2025). 10.1038/s41588-025-02314-y

18 Xin, B. et al. Colon cancer secreted protein-2 (CCSP-2), a novel candidate serological marker of colon neoplasia. Oncogene 24, 724–731 (2005). 10.1038/sj.onc.1208134

19 Lu, L., Johnson, C. H., Khan, S. A. & Irwin, M. L. GDF15 in the tumor microenvironment: A central mediator of cancer immunometabolism and therapeutic resistance. Cytokine & Growth Factor Reviews 88, 47–57 (2026). 10.1016/j.cytogfr.2026.01.004

20 Aoki, K., Nitta, A. & Igarashi, A. CDX1 and CDX2 suppress colon cancer stemness by inhibiting β-catenin-facilitated formation of Pol II–DSIF–PAF1C complex. Cell Death & Disease 16, 408 (2025). 10.1038/s41419-025-07737-3

21 Faupel-Badger, J. et al. Defining precancer: a grand challenge for the cancer community. Nature Reviews. Cancer 24, 792–809 (2024). 10.1038/s41568-024-00744-0

22 Stangis, M. M. et al. The Hallmarks of Precancer. Cancer Discovery 14, 683–689 (2024). 10.1158/2159-8290.CD-23-1550

23 Staton, C. A. et al. The angiogenic switch occurs at the adenoma stage of the adenoma carcinoma sequence in colorectal cancer. Gut 56, 1426–1432 (2007). 10.1136/gut.2007.125286

24 Guo, Y. et al. Senescence-associated tissue microenvironment promotes colon cancer formation through the secretory factor GDF15. Aging Cell 18, e13013 (2019). 10.1111/acel.13013

25 Ritschka, B. et al. The senescence-associated secretory phenotype induces cellular plasticity and tissue regeneration. Genes & Development 31, 172–183 (2017). 10.1101/gad.290635.116

26 De Blander, H., Morel, A.-P., Senaratne, A. P., Ouzounova, M. & Puisieux, A. Cellular Plasticity: A Route to Senescence Exit and Tumorigenesis. Cancers 13, 4561 (2021). 10.3390/cancers13184561

27 Wang, D. et al. GDF15: emerging biology and therapeutic applications for obesity and cardiometabolic disease. Nat Rev Endocrinol 17, 592–607 (2021). 10.1038/s41574-021-00529-7

28 Wischhusen, J., Melero, I. & Fridman, W. H. Growth/Differentiation Factor-15 (GDF-15): From Biomarker to Novel Targetable Immune Checkpoint. Frontiers in Immunology 11 (2020). 10.3389/fimmu.2020.00951

29 Haake, M. et al. Tumor-derived GDF-15 blocks LFA-1 dependent T cell recruitment and suppresses responses to anti-PD-1 treatment. Nature Communications 14, 4253 (2023). 10.1038/s41467-023-39817-3

30 Guan, B. et al. Peritumoral colonic epithelial cell-derived GDF15 sustains colorectal cancer via regulation of glycolysis and histone lactylation. Nature Aging 5, 2449–2465 (2025). 10.1038/s43587-025-01023-9

31 He, X. et al. Long-term Risk of Colorectal Cancer After Removal of Conventional Adenomas and Serrated Polyps. Gastroenterology 158, 852–861 e854 (2020). 10.1053/j.gastro.2019.06.039

32 Rey, S. J. & Anselin, L. PySAL: A Python Library of Spatial Analytical Methods. Review of Regional Studies 37 (2007). 10.52324/001c.8285

33 Marconato, L. et al. SpatialData: an open and universal data framework for spatial omics. Nature Methods 22, 58–62 (2025). 10.1038/s41592-024-02212-x

34 Pedregosa, F. et al. Scikit-learn: Machine Learning in Python. Journal of Machine Learning Research 12, 2825–2830 (2011).

35 Virshup, I., Rybakov, S., Theis, F. J., Angerer, P. & Wolf, F. A. anndata: Access and store annotated data matrices. Journal of Open Source Software 9, 4371 (2024). 10.21105/joss.04371

36 Wolf, F. A., Angerer, P. & Theis, F. J. SCANPY: large-scale single-cell gene expression data analysis. Genome Biology 19, 15 (2018). 10.1186/s13059-017-1382-0

37 Korsunsky, I. et al. Fast, sensitive and accurate integration of single-cell data with Harmony. Nature Methods 16, 1289–1296 (2019). 10.1038/s41592-019-0619-0

38 Palla, G. et al. Squidpy: a scalable framework for spatial omics analysis. Nature Methods 19, 171–178 (2022). 10.1038/s41592-021-01358-2

39 Seabold, S. & Perktold, J. Statsmodels: Econometric and Statistical Modeling with Python. SciPy 2010 (2010). 10.25080/Majora-92bf1922-011

