## Supplementary Figures for "Spatial multi-omics and single-cell transcriptomics uncover senescence-associated cellular programs during colon adenoma to cancer progression"

**Supplementary Information**

**Supplementary Figures**

**Supplementary Figure 1.** Overview of spatial multi-omic data processing, integration, and annotation.

**Supplementary Figure 2.** Spatial maps of all Visium adenoma samples.

**Supplementary Figure 3.** Characterization and annotation of Visium spatial clusters.

**Supplementary Figure 4.** Single-cell reference-guided spatial mapping reveals senescence-stemness colocalization and CD8 T-cell exclusion in adenomas.

**Supplementary Figure 5.** Cell2location-inferred cell-type abundance across spatial spots.

**Supplementary Figure 6.** Independent scRNA-seq datasets map senescence and stemness programs to distinct dysplastic epithelial subpopulations in adenomas.

**Supplementary Figure 7.** GDF15 marks dysplastic epithelial states associated with senescence-stemness programs and spatial CD8 T-cell exclusion.

**Supplementary Figure 8.** Principal component analysis of Xenium tissue microarray (TMA) cores.

**Supplementary Figure 9.** GDF15 is predominantly expressed in epithelial cells.

**Supplementary Figure 10.** Xenium cell clustering and subclustering of non-epithelial cells.

**Supplementary Figure 11.** Characterization of Xenium epithelial clusters across patients and tissue types.

**Supplementary Figure 12.** Spatial patterns and markers of stemness, senescence, and GDF15 hotspots across tissue types.

**Supplementary Tables**

**Supplementary Table 1.** Clinical and histopathological features of the Visium adenoma cohort.

**Supplementary Table 2.** Clinical and histopathological features of the Xenium TMA cohort.

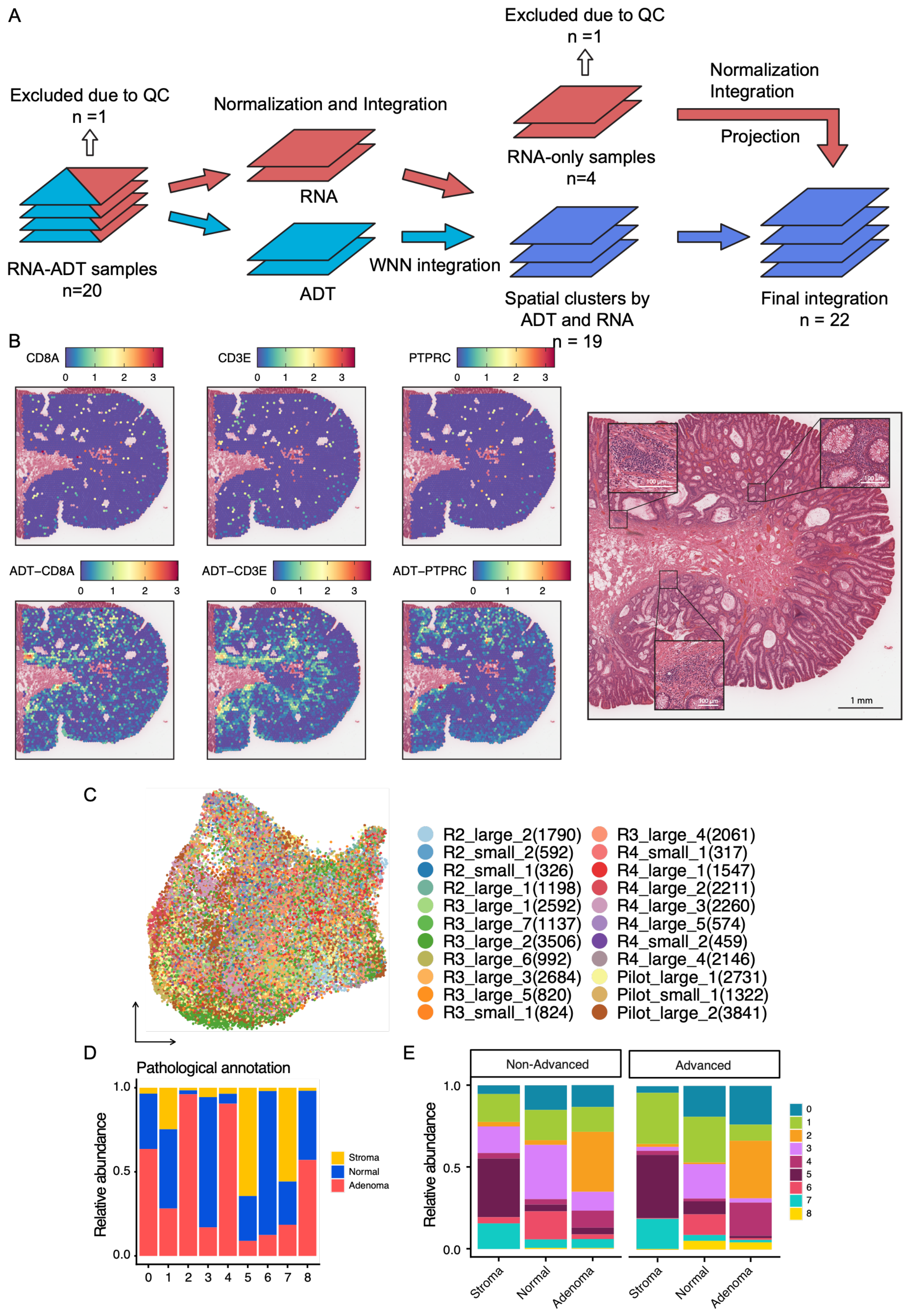
A
Excluded due to QC
n =1
RNA-ADT samples
n=20
Normalization and Integration
RNA
ADT
WNN integration
Excluded due to QC
n =1
RNA-only samples
n=4
Normalization
Integration
Projection
Spatial clusters by
ADT and RNA
n = 19
Final integration
n = 22
B
CD8A
CD3E
PTPRC
ADT−CD8A
ADT−CD3E
ADT−PTPRC
100 μm
1 mm
C
R2_large_2(1790)
R2_small_2(592)
R2_small_1(326)
R2_large_1(1198)
R3_large_1(2592)
R3_large_7(1137)
R3_large_2(3506)
R3_large_6(992)
R3_large_3(2684)
R3_large_5(820)
R3_small_1(824)
R3_large_4(2061)
R4_small_1(317)
R4_large_1(1547)
R4_large_2(2211)
R4_large_3(2260)
R4_large_5(574)
R4_small_2(459)
R4_large_4(2146)
Pilot_large_1(2731)
Pilot_small_1(1322)
Pilot_large_2(3841)
D
Pathological annotation
Relative abundance
Stroma
Normal
Adenoma
E
Non-Advanced
Advanced
Relative abundance
Stroma
Normal
Adenoma

Supplementary Figure 1. Overview of spatial multi-omic data processing, integration, and annotation.

(A) Schematic of the workflow used to preprocess and integrate Visium RNA and antibody-derived tag (ADT) data. The modalities were processed separately, dimensionally reduced, and integrated to generate a shared multimodal representation for clustering and annotation.

(B) Representative spatial feature plots of transcriptomic and protein-derived signals across tissue sections, shown with the corresponding H&E-stained sections and selected regions of interest.

(C) UMAP visualization of integrated spatial spots colored by sample. The broad intermixing of spots from different samples indicates minimal residual sample-level batch effects after integration.

(D) Stacked bar plot showing the distribution of pathologist-defined histologic annotations within each unsupervised spatial cluster.

(E) Stacked bar plots showing the spatial-cluster composition of each histologic annotation category, stratified by adenoma size group.

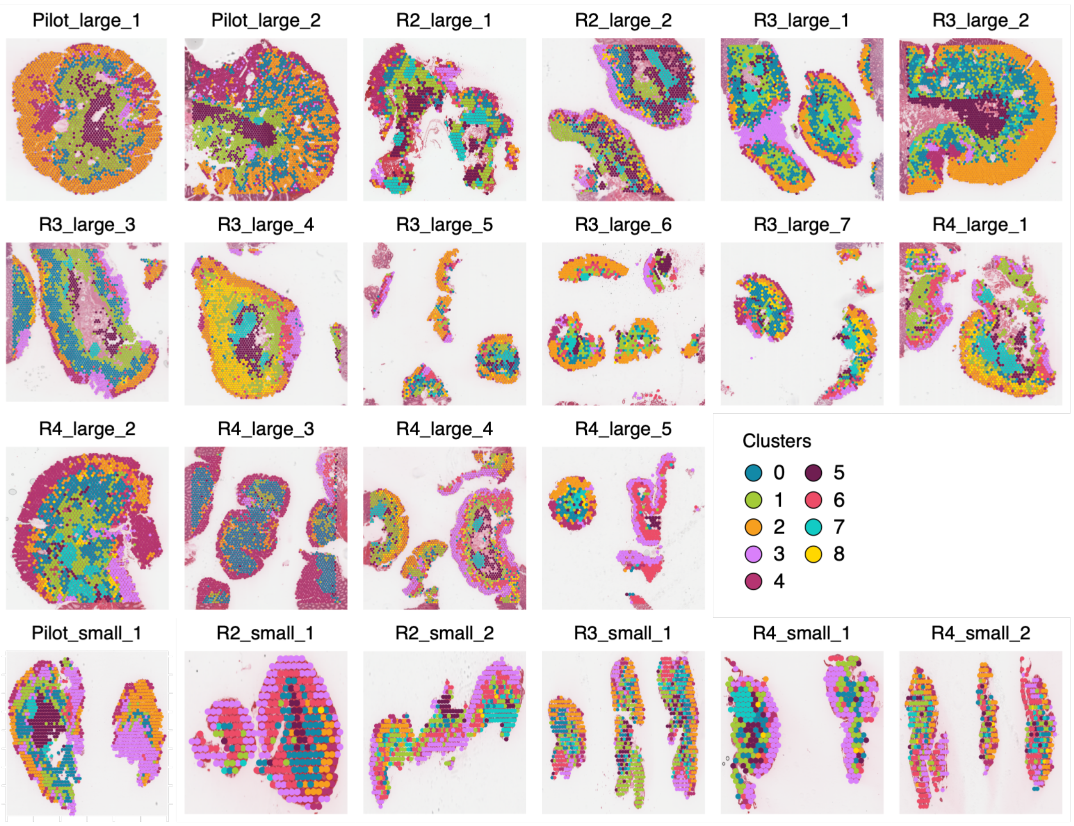

Pilot_large_1
Pilot_large_2
R2_large_1
R2_large_2
R3_large_1
R3_large_2
R3_large_3
R3_large_4
R3_large_5
R3_large_6
R3_large_7
R4_large_1
R4_large_2
R4_large_3
R4_large_4
R4_large_5
Clusters
0
1
2
3
4
5
6
7
8
Pilot_small_1
R2_small_1
R2_small_2
R3_small_1
R4_small_1
R4_small_2


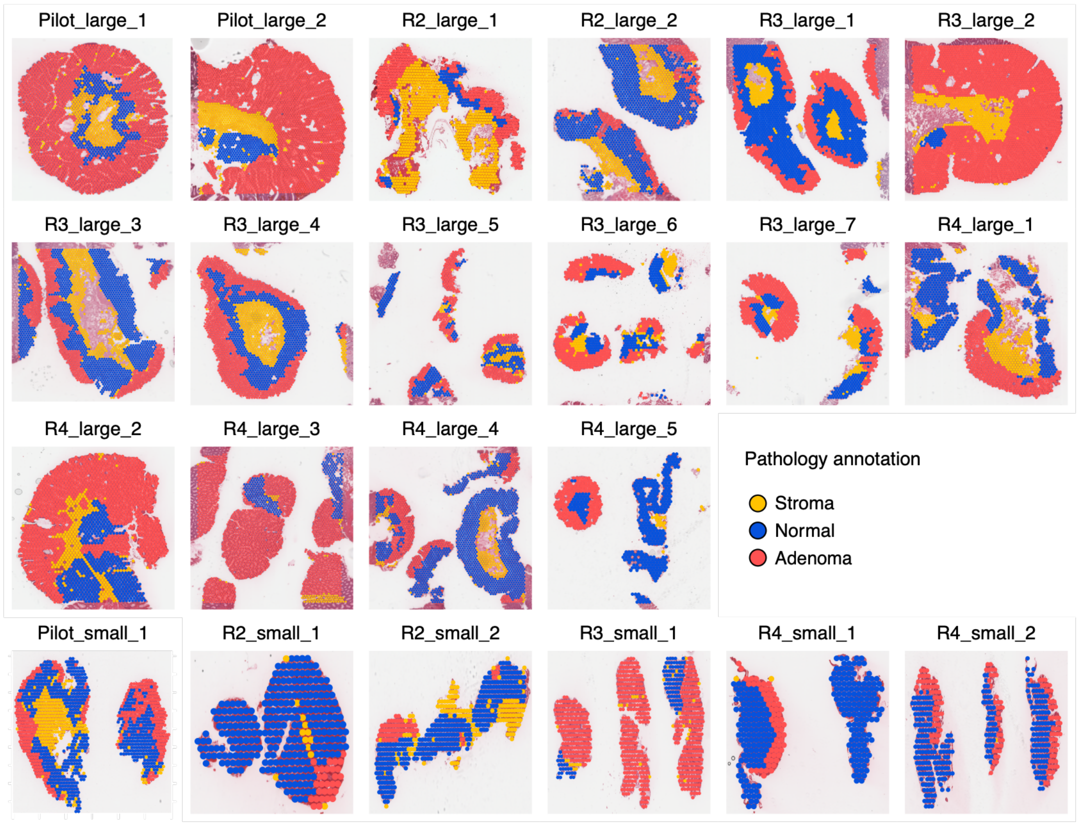

Pilot_large_1
Pilot_large_2
R2_large_1
R2_large_2
R3_large_1
R3_large_2
R3_large_3
R3_large_4
R3_large_5
R3_large_6
R3_large_7
R4_large_1
R4_large_2
R4_large_3
R4_large_4
R4_large_5
Pathology annotation
Stroma
Normal
Adenoma
Pilot_small_1
R2_small_1
R2_small_2
R3_small_1
R4_small_1
R4_small_2

Supplementary Figure 2. Spatial maps of all Visium adenoma samples.

(A) Spatial distribution of unsupervised clusters across all adenoma samples.

(B) Spatial distribution of pathologist-defined histologic annotations across all adenoma samples.

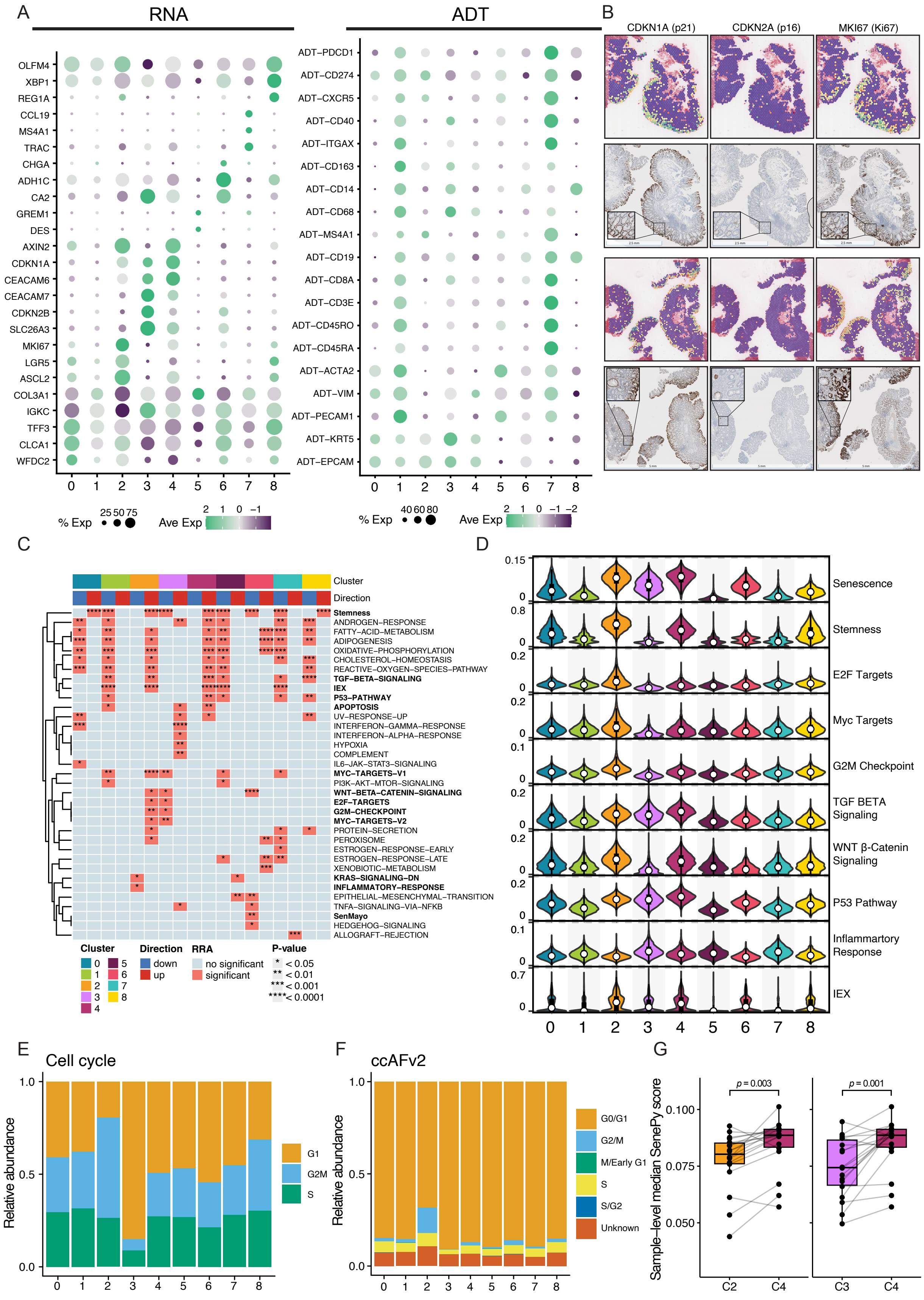

A
RNA
ADT
OLFM4
XBP1
REG1A
CCL19
MS4A1
TRAC
CHGA
ADH1C
CA2
GREM1
DES
AXIN2
CDKN1A
CEACAM6
CEACAM7
CDKN2B
SLC26A3
MKI67
LGR5
ASCL2
COL3A1
IGKC
TFF3
CLCA1
WFDC2
0 1 2 3 4 5 6 7 8
% Exp 25 50 75
Ave Exp 2 1 0 −1
ADT−PDCD1
ADT−CD274
ADT−CXCR5
ADT−CD40
ADT−ITGAX
ADT−CD163
ADT−CD14
ADT−CD68
ADT−MS4A1
ADT−CD19
ADT−CD8A
ADT−CD3E
ADT−CD45RO
ADT−CD45RA
ADT−ACTA2
ADT−VIM
ADT−PECAM1
ADT−KRT5
ADT−EPCAM
0 1 2 3 4 5 6 7 8
% Exp 40 60 80
Ave Exp 2 1 0 −1 −2
B
CDKN1A (p21)
CDKN2A (p16)
MKI67 (Ki67)
C
Cluster
Direction
Stemness
ANDROGEN−RESPONSE
FATTY−ACID−METABOLISM
ADIPOGENESIS
OXIDATIVE−PHOSPHORYLATION
CHOLESTEROL−HOMEOSTASIS
REACTIVE−OXYGEN−SPECIES−PATHWAY
TGF−BETA−SIGNALING
IEX
P53−PATHWAY
APOPTOSIS
UV−RESPONSE−UP
INTERFERON−GAMMA−RESPONSE
INTERFERON−ALPHA−RESPONSE
HYPOXIA
COMPLEMENT
IL6−JAK−STAT3−SIGNALING
MYC−TARGETS−V1
PI3K−AKT−MTOR−SIGNALING
WNT−BETA−CATENIN−SIGNALING
E2F−TARGETS
G2M−CHECKPOINT
MYC−TARGETS−V2
PROTEIN−SECRETION
PEROXISOME
ESTROGEN−RESPONSE−EARLY
ESTROGEN−RESPONSE−LATE
XENOBIOTIC−METABOLISM
KRAS−SIGNALING−DN
INFLAMMATORY−RESPONSE
EPITHELIAL−MESENCHYMAL−TRANSITION
TNFA−SIGNALING−VIA−NFKB
SenMayo
HEDGEHOG−SIGNALING
ALLOGRAFT−REJECTION
Cluster 0 1 2 3 4 5 6 7 8
Direction down up
RRA no significant significant
P-value
* < 0.05
** < 0.01
*** < 0.001
**** < 0.0001
D
Senescence
Stemness
E2F Targets
Myc Targets
G2M Checkpoint
TGF BETA Signaling
WNT β-Catenin Signaling
P53 Pathway
Inflammartory Response
IEX
0 1 2 3 4 5 6 7 8
E
Cell cycle
Relative abundance
G1
G2M
S
0 1 2 3 4 5 6 7 8
F
ccAFv2
Relative abundance
G0/G1
G2/M
M/Early G1
S
S/G2
Unknown
0 1 2 3 4 5 6 7 8
G
Sample−level median SenePy score
p = 0.003
p = 0.001
C2 C4
C3 C4

Supplementary Figure 3. Characterization and annotation of Visium spatial clusters.

(A) Dot plots showing selected marker features measured by RNA (left) and antibody-derived tags (ADT; right) across spatial clusters. Dot color indicates scaled mean expression, and dot size indicates the percentage of spots expressing each feature.

(B) Representative spatial expression maps and immunohistochemical staining of consecutive tissue sections for CDKN1A (p21), CDKN2A (p16), and MKI67 (Ki-67). Spatial transcript levels are shown as log2-normalized counts on a scale from 0 to 3.

(C) Heatmap of gene set enrichment results integrated across scoring methods using Robust Rank Aggregation (RRA). Gene sets significantly enriched in at least one cluster are shown; those significant in clusters 2 and 4 are highlighted in bold. Asterisks indicate statistical significance.

(D) Stacked violin plots showing gene set activity scores across clusters. UCell scores are shown for all gene sets except senescence, which was scored using the SenePy universal hub module.

(E) Stacked bar plot showing cell-cycle phase assignments within each cluster based on Seurat cell-cycle scoring.

(F) Stacked bar plot showing cell-cycle phase assignments within each cluster inferred using ccAFv2.

(G) Box plots showing sample-level median SenePy universal hub scores in clusters 2, 3, and 4. Dots represent individual samples, and lines connect paired samples. Clusters 2 versus 4 and 3 versus 4 were compared using paired Wilcoxon signed-rank tests.

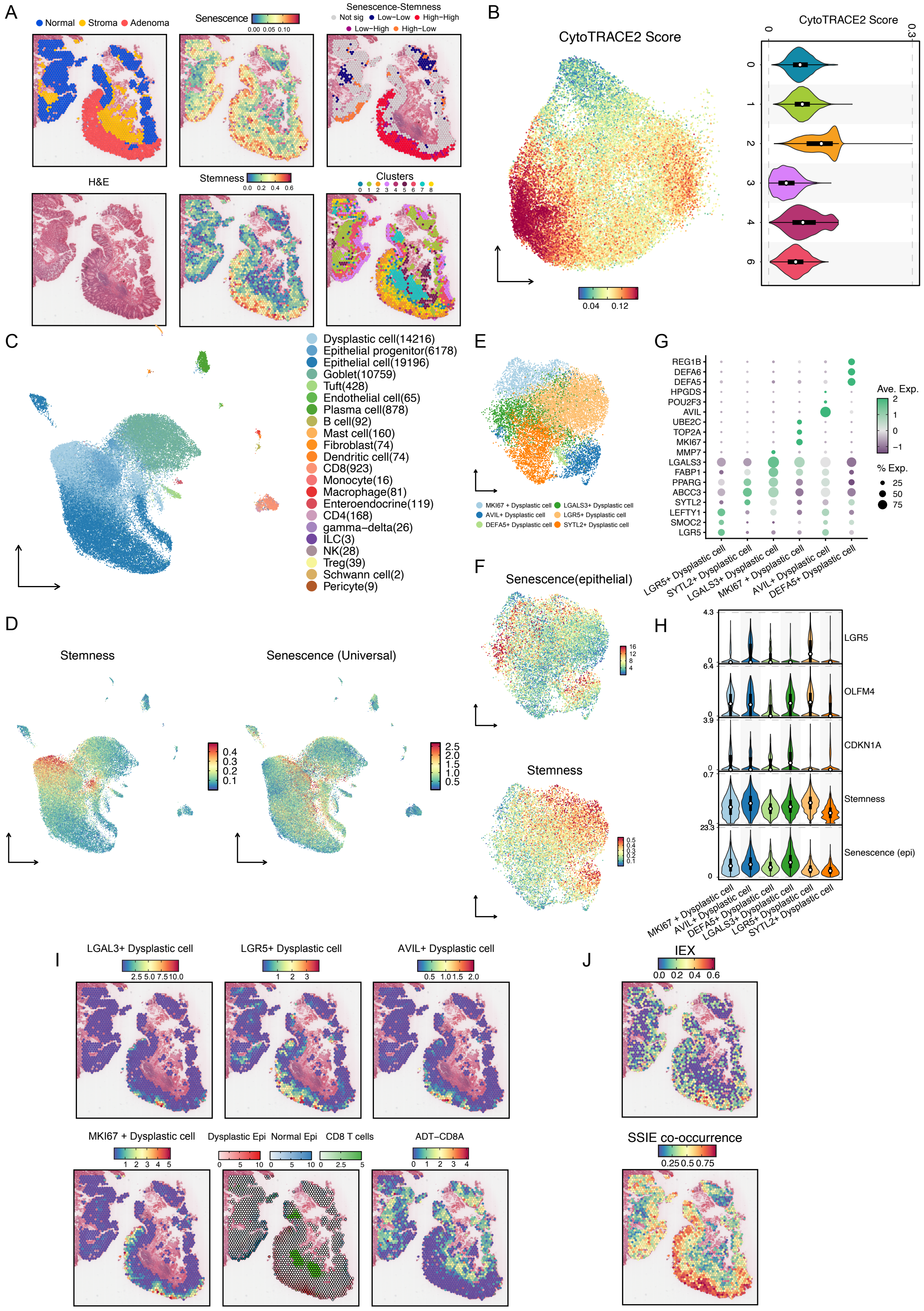

A
Normal
Stroma
Adenoma
Senescence
Senescence-Stemness
Not sig
Low−Low
High−High
Low−High
High−Low
H&E
Stemness
Clusters
B
CytoTRACE2 Score
C
Dysplastic cell(14216)
Epithelial progenitor(6178)
Epithelial cell(19196)
Goblet(10759)
Tuft(428)
Endothelial cell(65)
Plasma cell(878)
B cell(92)
Mast cell(160)
Fibroblast(74)
Dendritic cell(74)
CD8(923)
Monocyte(16)
Macrophage(81)
Enteroendocrine(119)
CD4(168)
gamma−delta(26)
ILC(3)
NK(28)
Treg(39)
Schwann cell(2)
Pericyte(9)
D
Stemness
Senescence (Universal)
E
MKI67 + Dysplastic cell
LGALS3+ Dysplastic cell
AVIL+ Dysplastic cell
LGR5+ Dysplastic cell
DEFA5+ Dysplastic cell
SYTL2+ Dysplastic cell
F
Senescence(epithelial)
Stemness
G
REG1B
DEFA6
DEFA5
HPGDS
POU2F3
AVIL
UBE2C
TOP2A
MKI67
MMP7
LGALS3
FABP1
PPARG
ABCC3
SYTL2
LEFTY1
SMOC2
LGR5
Ave. Exp.
% Exp.
LGR5+ Dysplastic cell
SYTL2+ Dysplastic cell
LGALS3+ Dysplastic cell
MKI67 + Dysplastic cell
AVIL+ Dysplastic cell
DEFA5+ Dysplastic cell
H
LGR5
OLFM4
CDKN1A
Stemness
Senescence (epi)
MKI67 + Dysplastic cell
AVIL+ Dysplastic cell
DEFA5+ Dysplastic cell
LGALS3+ Dysplastic cell
LGR5+ Dysplastic cell
SYTL2+ Dysplastic cell
I
LGAL3+ Dysplastic cell
LGR5+ Dysplastic cell
AVIL+ Dysplastic cell
MKI67 + Dysplastic cell
Dysplastic Epi
Normal Epi
CD8 T cells
ADT−CD8A
J
IEX
SSIE co-occurrence

Supplementary Figure 4. Single-cell reference-guided spatial mapping reveals senescence-stemness colocalization and CD8 T-cell exclusion in adenomas.

(A) Representative spatial maps of a large adenoma showing histologic annotations, H&E staining, senescence and stemness scores, local bivariate Moran's I classes for senescence-stemness associations, and spatial cluster assignments.

(B) Spatial UMAP visualization and violin plot showing CytoTRACE2 scores across epithelial spatial clusters; higher scores indicate greater developmental potential, a proxy for stemness.

(C) UMAP visualization of the curated public scRNA-seq dataset used as the deconvolution reference, colored by major cell type.

(D) UMAP visualizations of stemness and universal senescence signature scores in the scRNA-seq reference dataset.

(E) UMAP visualization of dysplastic epithelial subtypes identified in the curated public scRNA-seq reference dataset.

(F) UMAP visualizations of epithelial senescence and stemness signature scores across dysplastic epithelial cells in the scRNA-seq reference dataset.

(G) Dot plot showing representative marker genes used to annotate dysplastic epithelial subtypes in the scRNA-seq reference. Dot color indicates scaled mean expression, and dot size indicates the percentage of cells expressing each gene.

(H) Violin plots showing representative stemness- and senescence-associated genes and signature scores across dysplastic epithelial subtypes.

(I) Representative spatial maps showing the deconvolution-inferred abundance of dysplastic epithelial subtypes, normal epithelial cells, and CD8 T cells, together with spatial ADT-CD8A protein signal.

(J) Representative spatial maps showing the Immune exclusion signature (IEX) and the Senescence–Stemness–Immune Exclusion Co-occurrence Score (SSIE).

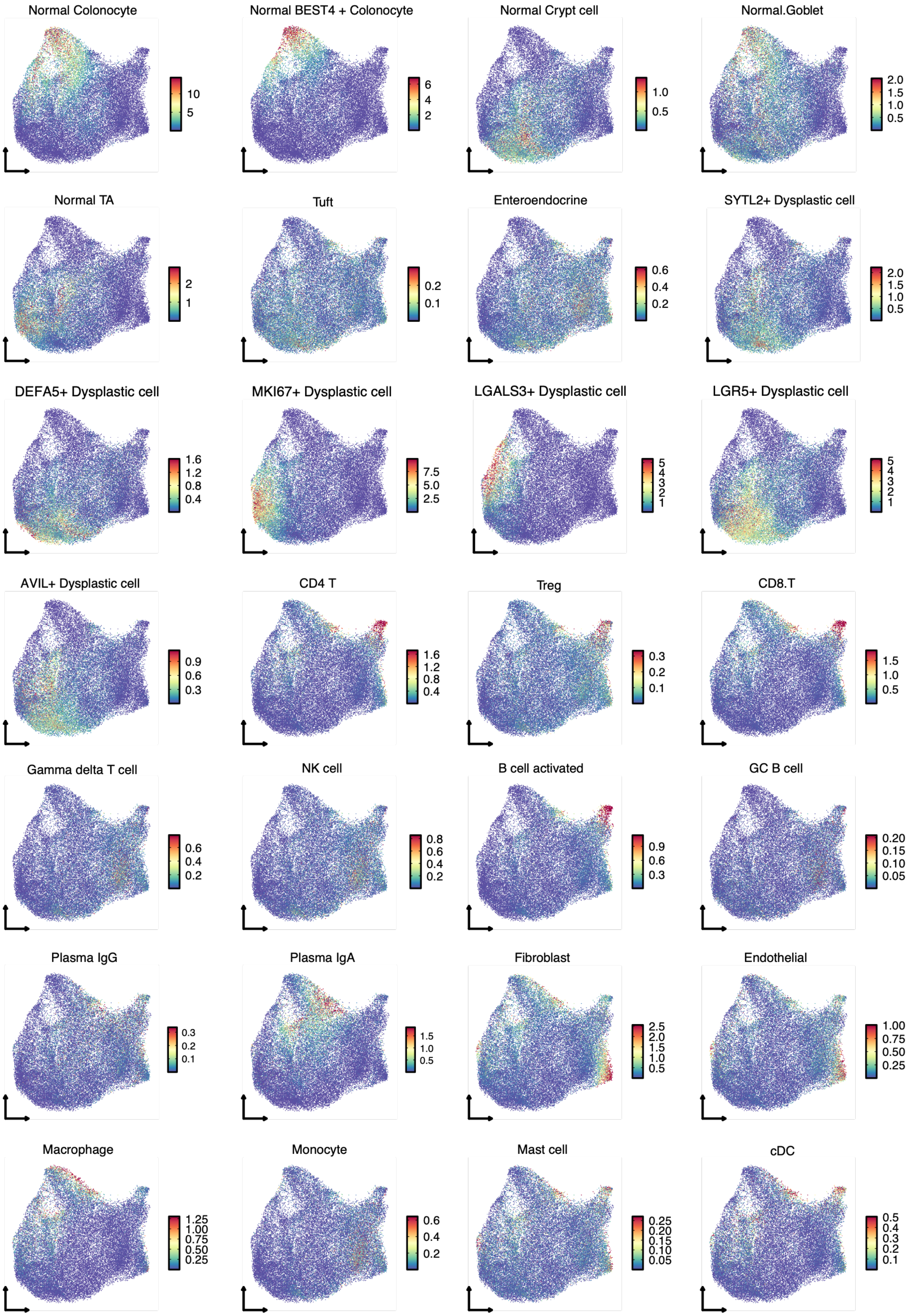
Normal Colonocyte
10
5
Normal BEST4 + Colonocyte
6
4
2
Normal Crypt cell
1.0
0.5
Normal.Goblet
2.0
1.5
1.0
0.5
Normal TA
2
1
Tuft
0.2
0.1
Enteroendocrine
0.6
0.4
0.2
SYTL2+ Dysplastic cell
2.0
1.5
1.0
0.5
DEFA5+ Dysplastic cell
1.6
1.2
0.8
0.4
MKI67+ Dysplastic cell
7.5
5.0
2.5
LGALS3+ Dysplastic cell
5
4
3
2
1
LGR5+ Dysplastic cell
5
4
3
2
1
AVIL+ Dysplastic cell
0.9
0.6
0.3
CD4 T
1.6
1.2
0.8
0.4
Treg
0.3
0.2
0.1
CD8.T
1.5
1.0
0.5
Gamma delta T cell
0.6
0.4
0.2
NK cell
0.8
0.6
0.4
0.2
B cell activated
0.9
0.6
0.3
GC B cell
0.20
0.15
0.10
0.05
Plasma IgG
0.3
0.2
0.1
Plasma IgA
1.5
1.0
0.5
Fibroblast
2.5
2.0
1.5
1.0
0.5
Endothelial
1.00
0.75
0.50
0.25
Macrophage
1.25
1.00
0.75
0.50
0.25
Monocyte
0.6
0.4
0.2
Mast cell
0.25
0.20
0.15
0.10
0.05
cDC
0.5
0.4
0.3
0.2
0.1

Supplementary Figure 5. Cell2location-inferred cell-type abundance across spatial spots.

UMAP visualization showing Cell2location-inferred abundance of selected cell types across Visium spots from all adenoma samples.

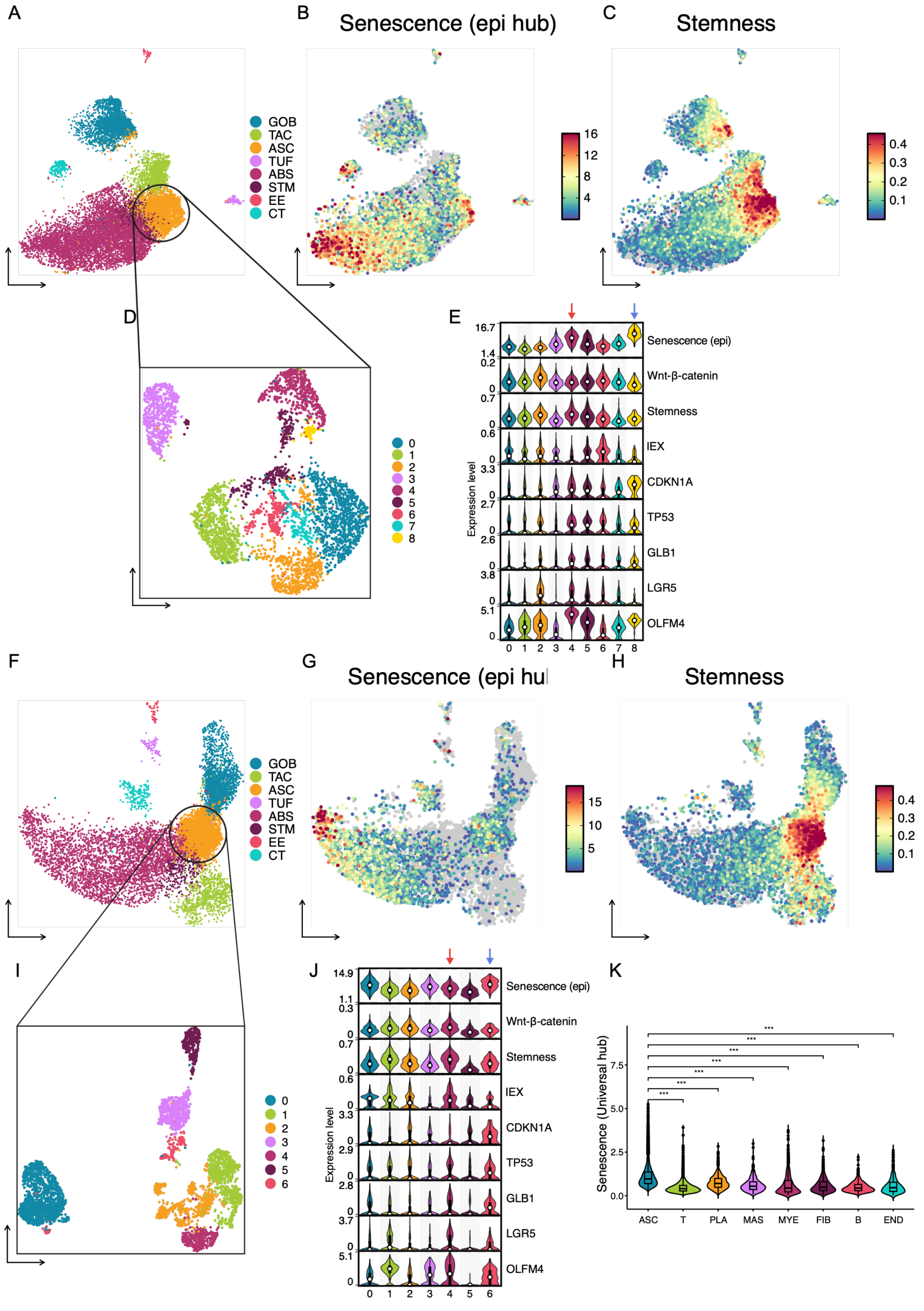

A
B
Senescence (epi hub)
C
Stemness
GOB
TAC
ASC
TUF
ABS
STM
EE
CT
D
E
Senescence (epi)
Wnt-β-catenin
Stemness
IEX
CDKN1A
TP53
GLB1
LGR5
OLFM4
Expression level
F
G
Senescence (epi hu
H
Stemness
I
J
K
Senescence (Universal hub)
ASC
T
PLA
MAS
MYE
FIB
B
END

Supplementary Figure 6. Independent scRNA-seq datasets map senescence and stemness programs to distinct dysplastic epithelial subpopulations in adenomas.

(A) UMAP visualization of epithelial cell populations from adenoma scRNA-seq dataset A, reported by Chen et al. (2021). Cell identities include goblet cells (GOB), transit-amplifying cells (TAC), adenoma-specific cells (ASC), tuft cells (TUF), absorptive cells (ABS), stem cells (STM), enteroendocrine cells (EE), and cycling/transit cells (CT).

(B) UMAP visualization of SenePy senescence scores in dataset A.

(C) UMAP visualization of stemness scores in dataset A, computed using the curated intestinal stemness signature applied in the Visium analyses.

(D) UMAP visualization of reclustered adenoma-specific epithelial cells from dataset A.

(E) Violin plots showing SenePy senescence scores, Wnt/β-catenin scores, stemness scores, immune exclusion (IEX) scores, and expression of selected marker genes across adenoma-specific epithelial subclusters in dataset A. Red arrows mark subclusters with the highest stemness scores, and blue arrows mark those with the highest senescence scores.

(F) UMAP visualization of epithelial cell populations in adenoma scRNA-seq dataset B from the same study.

(G) UMAP visualization of SenePy senescence scores in dataset B.

(H) UMAP visualization of stemness scores in dataset B, computed using the curated intestinal stemness signature applied in the Visium analyses.

(I) UMAP visualization of reclustered adenoma-specific epithelial cells from dataset B.

(J) Violin plots showing SenePy senescence scores, Wnt/β-catenin scores, stemness scores, IEX scores, and expression of selected marker genes across adenoma-specific epithelial subclusters in dataset B. Red arrows mark subclusters with the highest stemness scores, and blue arrows mark those with the highest senescence scores.

(K) Integrated analysis of epithelial datasets A and B and the non-epithelial scRNA-seq dataset, showing SenePy universal scores across major cell types. ASC, adenoma-specific cells; PLA, plasma cells; MAS, mast cells; MYE, myeloid cells; FIB, fibroblasts; END, endothelial cells. Asterisks indicate statistical significance. *** $P < 0.001$.

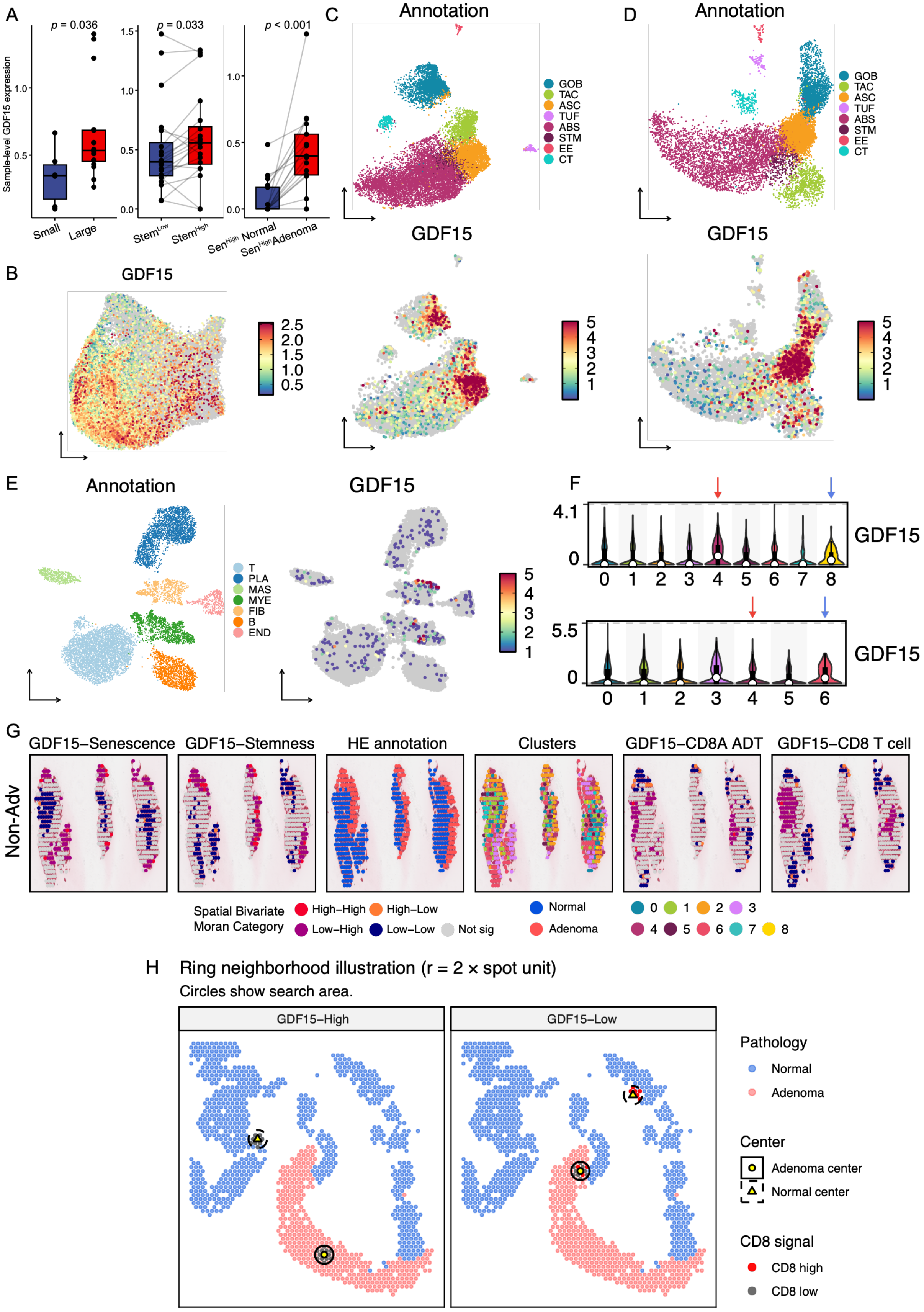
A
p = 0.036
p = 0.033
p < 0.001
Sample-level GDF15 expression
Small
Large
Stem^Low
Stem^High
Sen^High Normal
Sen^High Adenoma
B
GDF15
C
Annotation
GOB
TAC
ASC
TUF
ABS
STM
EE
CT
GDF15
D
Annotation
GDF15
E
Annotation
T
PLA
MAS
MYE
FIB
B
END
GDF15
F
GDF15
G
GDF15−Senescence
GDF15−Stemness
HE annotation
Clusters
GDF15−CD8A ADT
GDF15−CD8 T cell
Non-Adv
Spatial Bivariate Moran Category
High−High
High−Low
Low−High
Low−Low
Not sig
Normal
Adenoma
H
Ring neighborhood illustration (r = 2 × spot unit)
Circles show search area.
GDF15−High
GDF15−Low
Pathology
Normal
Adenoma
Center
Adenoma center
Normal center
CD8 signal
CD8 high
CD8 low

Supplementary Figure 7. GDF15 marks dysplastic epithelial states associated with senescence-stemness programs and spatial CD8 T-cell exclusion.

(A) Box plots comparing sample-level median GDF15 expression in small versus large adenomas (left), in stemness-low versus stemness-high spots (middle), and in senescence-high spots from normal versus adenoma regions (right). Dots represent individual samples; lines connect paired samples in the middle and right comparisons. The small-versus-large comparison used a Wilcoxon rank-sum test; the middle and right comparisons used paired Wilcoxon signed-rank tests.

(B) UMAP visualization showing GDF15 expression across Visium spots from all adenoma samples.

(C) UMAP visualizations of epithelial cell populations and GDF15 expression in public adenoma scRNA-seq dataset A.

(D) UMAP visualizations of epithelial cell populations and GDF15 expression in dataset B from the same study.

(E) UMAP visualizations of non-epithelial cell populations and GDF15 expression in the public scRNA-seq dataset.

(F) Violin plots showing GDF15 expression across adenoma-specific cell (ASC) subclusters in public epithelial scRNA-seq datasets A (top) and B (bottom). Red arrows mark subclusters with the highest stemness scores, and blue arrows mark those with the highest senescence scores.

(G) Representative spatial maps of a small adenoma showing bivariate spatial associations between GDF15 and senescence, GDF15 and stemness, GDF15 and ADT-CD8A, and GDF15 and inferred CD8 T-cell abundance, together with H&E-based histologic annotations and spatial clusters. Associations were classified using bivariate local Moran's I.

(H) Schematic of the radius-based spatial neighborhood analysis used to quantify CD8 T-cell abundance around GDF15-high and GDF15-low centers in normal and adenoma regions. Circles denote a search radius of two Visium spot units. ASC, adenoma-specific cells; GOB, goblet cells; TAC, transit-amplifying cells; TUF, tuft cells; ABS, absorptive cells; STM, stem cells; EE, enteroendocrine cells; CT, cycling/transit cells; PLA, plasma cells; MAS, mast cells; MYE, myeloid cells; FIB, fibroblasts; END, endothelial cells.

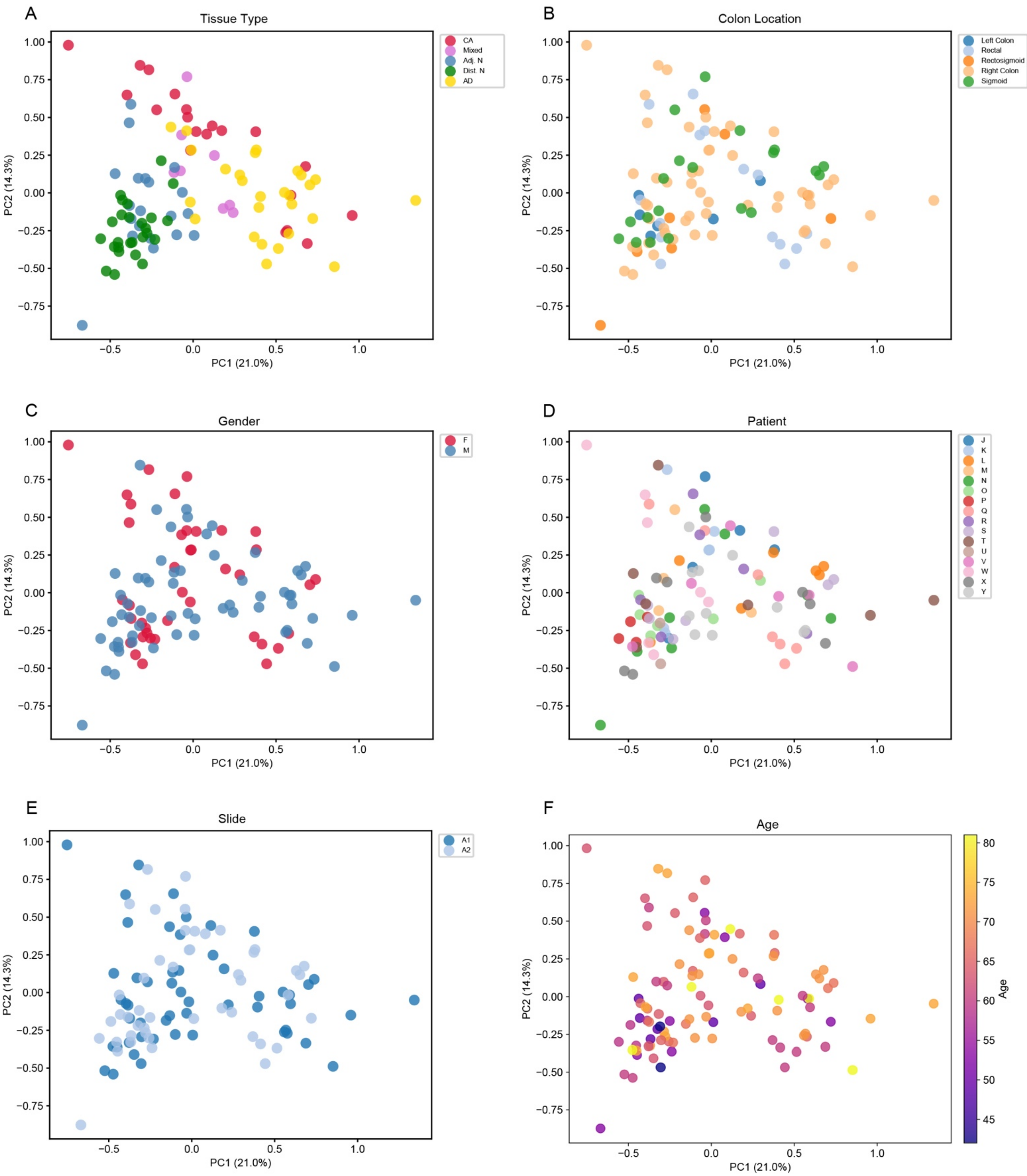


Supplementary Figure 8. Principal component analysis of Xenium tissue microarray (TMA) cores.

Principal component analysis (PCA) of all Xenium TMA cores using 10 principal components (PCs). Cores are colored by (A) tissue type, (B) colon location, (C) gender, (D) patient, (E) TMA slide, and (F) age. AD, conventional adenoma; CA, carcinoma; Adj. N, adjacent normal; Dist. N, distant normal.

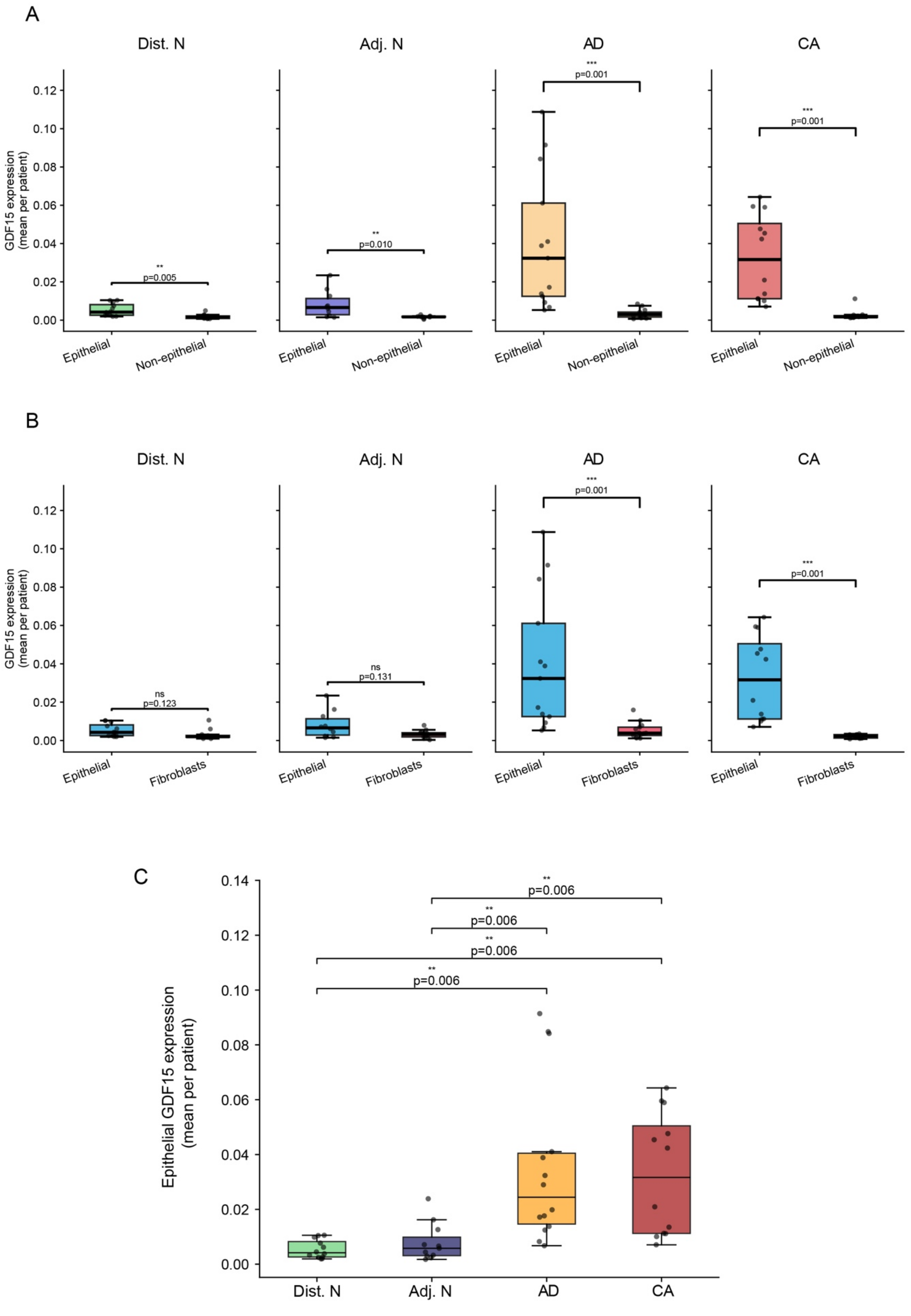

A
Dist. N
Adj. N
AD
CA
GDF15 expression (mean per patient)
0.00
0.02
0.04
0.06
0.08
0.10
0.12
**
p=0.005
**
p=0.010
***
p=0.001
***
p=0.001
Epithelial
Non-epithelial
B
Dist. N
Adj. N
AD
CA
GDF15 expression (mean per patient)
ns
p=0.123
ns
p=0.131
***
p=0.001
***
p=0.001
Epithelial
Fibroblasts
C
Epithelial GDF15 expression (mean per patient)
0.00
0.02
0.04
0.06
0.08
0.10
0.12
0.14
**
p=0.006
**
p=0.006
**
p=0.006
**
p=0.006
Dist. N
Adj. N
AD
CA

Supplementary Figure 9. GDF15 is predominantly expressed in epithelial cells.

(A) Box plots comparing normalized GDF15 expression between epithelial and non-epithelial cells within each tissue type. Significance was assessed using two-sided Wilcoxon signed-rank tests paired by patient, with Benjamini-Hochberg false discovery rate correction for multiple comparisons. Mixed-tissue cores were excluded.
(B) Box plots comparing normalized GDF15 expression between epithelial cells and fibroblasts within each tissue type. Significance was assessed using two-sided Wilcoxon signed-rank tests paired by patient, with Benjamini-Hochberg false discovery rate correction for multiple comparisons. Mixed-tissue cores were excluded.
(C) Box plot comparing normalized GDF15 expression in epithelial cells across tissue types. Significance was assessed for six pairwise comparisons using two-sided Wilcoxon signed-rank tests paired by patient, with Benjamini-Hochberg false discovery rate correction for multiple comparisons. Mixed-tissue cores were excluded. AD, conventional adenoma; CA, carcinoma; Adj. N, adjacent normal; Dist. N, distant normal.

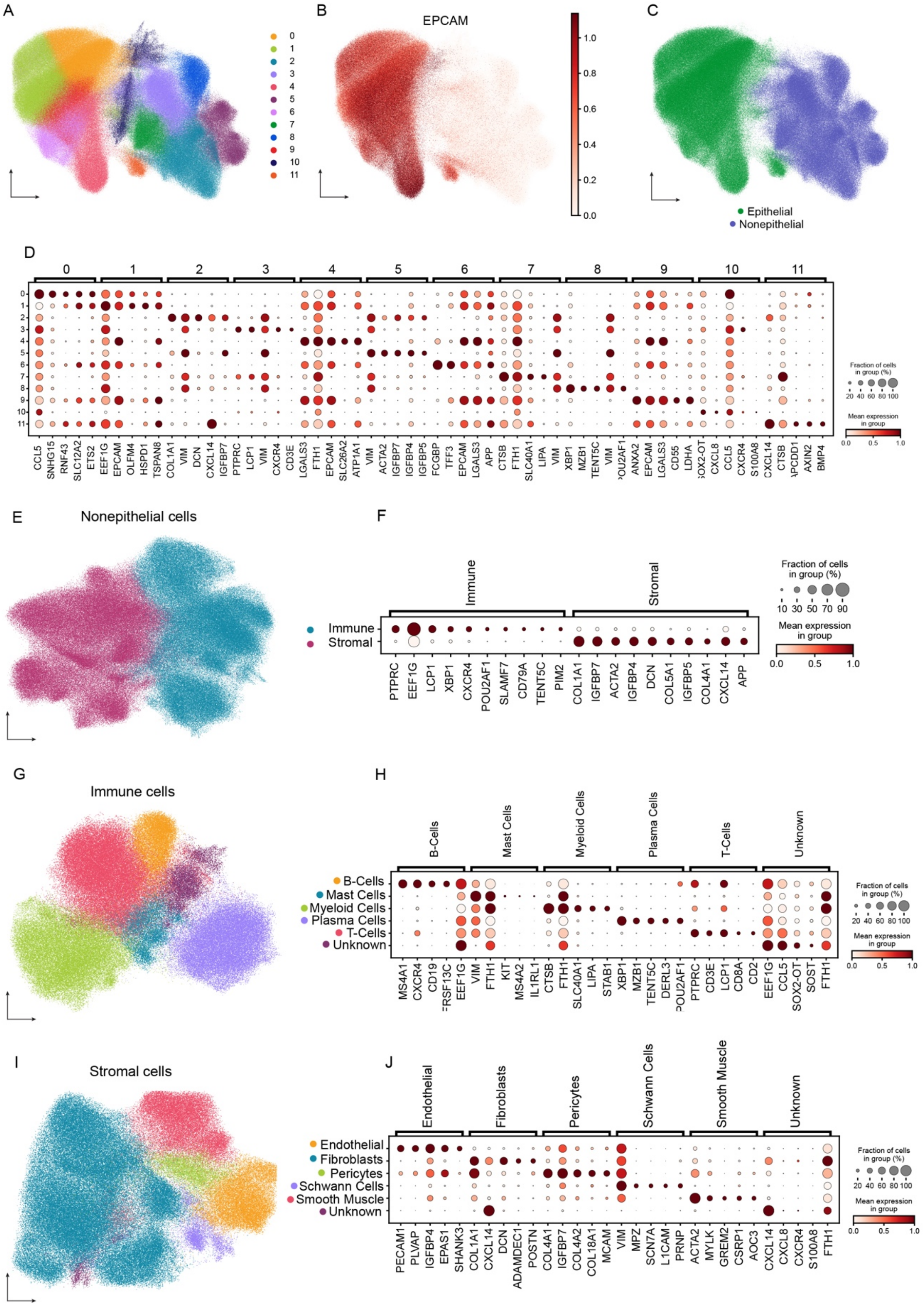

A
0
1
2
3
4
5
6
7
8
9
10
11
B
EPCAM
1.0
0.8
0.6
0.4
0.2
0.0
C
Epithelial
Nonepithelial
D
0
1
2
3
4
5
6
7
8
9
10
11
CCL5 SNHG15 RNF43 SLC12A2 ETS2 EEF1G EPCAM OLFM4 HSPD1 TSPAN8 COL1A1 VIM DCN CXCL14 IGFBP7 PTPRC LCP1 VIM CXCR4 CD3E LGALS3 FTH1 EPCAM SLC26A2 ATP1A1 VIM ACTA2 IGFBP7 IGFBP4 IGFBP5 FCGBP TFF3 EPCAM LGALS3 APP CTSB FTH1 SLC40A1 LIPA VIM XBP1 MZB1 TENT5C VIM POU2AF1 ANXA2 EPCAM LGALS3 CD55 LDHA SOX2-OT CXCL8 CCL5 CXCR4 S100A8 CXCL14 CTSB APCDD1 AXIN2 BMP4
Fraction of cells in group (%)
20 40 60 80 100
Mean expression in group
0.0 0.5 1.0
E
Nonepithelial cells
F
Immune
Stromal
Immune
Stromal
PTPRC EEF1G LCP1 XBP1 CXCR4 POU2AF1 SLAMF7 CD79A TENT5C PIM2 COL1A1 IGFBP7 ACTA2 IGFBP4 DCN COL5A1 IGFBP5 COL4A1 CXCL14 APP
Fraction of cells in group (%)
10 30 50 70 90
Mean expression in group
0.0 0.5 1.0
G
Immune cells
H
B-Cells
Mast Cells
Myeloid Cells
Plasma Cells
T-Cells
Unknown
B-Cells
Mast Cells
Myeloid Cells
Plasma Cells
T-Cells
Unknown
MS4A1 CXCR4 CD19 TNFRSF13C EEF1G VIM FTH1 KIT MS4A2 IL1RL1 CTSB FTH1 SLC40A1 LIPA STAB1 XBP1 MZB1 TENT5C DERL3 POU2AF1 PTPRC CD3E LCP1 CD8A CD2 EEF1G CCL5 SOX2-OT SOST FTH1
Fraction of cells in group (%)
20 40 60 80 100
Mean expression in group
0.0 0.5 1.0
I
Stromal cells
J
Endothelial
Fibroblasts
Pericytes
Schwann Cells
Smooth Muscle
Unknown
Endothelial
Fibroblasts
Pericytes
Schwann Cells
Smooth Muscle
Unknown
PECAM1 PLVAP IGFBP4 EPAS1 SHANK3 COL1A1 CXCL14 DCN ADAMDEC1 POSTN COL4A1 IGFBP7 COL4A2 COL18A1 MCAM VIM MPZ SCN7A L1CAM PRNP ACTA2 MYLK GREM2 CSRP1 AOC3 CXCL14 CXCL8 CXCR4 S100A8 FTH1
Fraction of cells in group (%)
20 40 60 80 100
Mean expression in group
0.0 0.5 1.0

Supplementary Figure 10. Xenium cell clustering and subclustering of non-epithelial cells.

(A) UMAP visualization of Louvain clusters from 101 Xenium cores across 16 patients, colored by cluster. Cluster 10 was excluded from downstream analyses.
(B) UMAP visualization colored by normalized EPCAM expression. Values were clipped at the 5th and 95th percentiles to limit the visual influence of outliers.
(C) UMAP visualization colored by epithelial and non-epithelial annotation after removal of cluster 10.
(D) Dot plot showing the top five marker genes for each cluster in (A). Dot size represents the percentage of cells expressing each gene, and color represents mean expression scaled within each gene.
(E) UMAP visualization of subclustered non-epithelial cells colored by broad immune and stromal annotation.
(F) Dot plot showing the top 10 marker genes for the immune and stromal clusters in (E). Dot size represents the percentage of cells expressing each gene, and color represents mean expression scaled within each gene.
(G) UMAP visualization of subclustered immune cells colored by annotated immune cell type.
(H) Dot plot showing the top five marker genes for each immune cluster in (G). Dot size represents the percentage of cells expressing each gene, and color represents mean expression scaled within each gene.
(I) UMAP visualization of subclustered stromal cells colored by annotated stromal cell type.
(J) Dot plot showing the top five marker genes for each stromal cluster in (I). Dot size represents the percentage of cells expressing each gene, and color represents mean expression scaled within each gene.

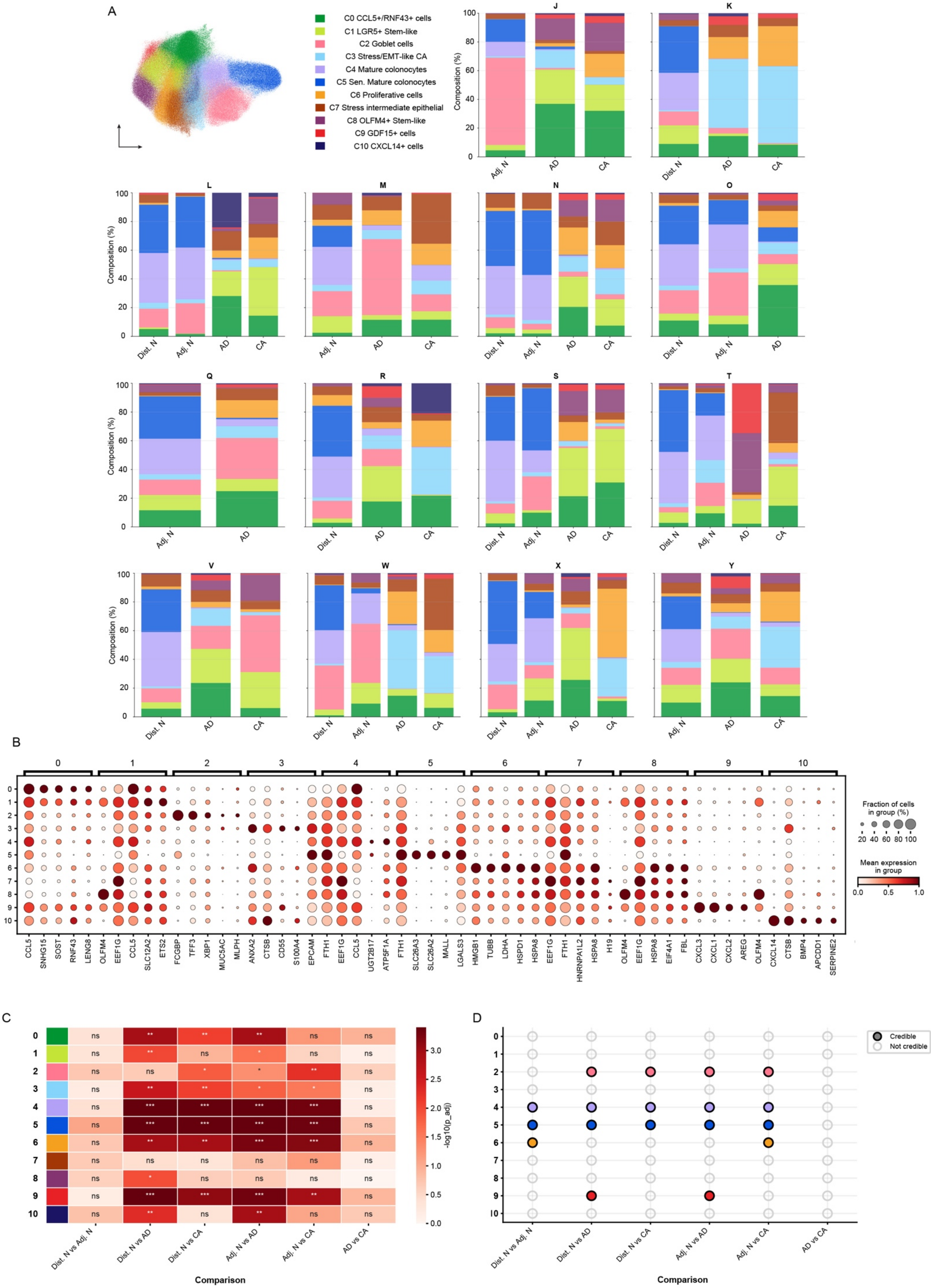
A
C0 CCL5+/RNF43+ cells
C1 LGR5+ Stem-like
C2 Goblet cells
C3 Stress/EMT-like CA
C4 Mature colonocytes
C5 Sen. Mature colonocytes
C6 Proliferative cells
C7 Stress intermediate epithelial
C8 OLFM4+ Stem-like
C9 GDF15+ cells
C10 CXCL14+ cells
J
K
L
M
N
O
Q
R
S
T
V
W
X
Y
Composition (%)
Dist. N
Adj. N
AD
CA
B
CCL5 SNHG15 SOST RNF43 LENG8 OLFM4 EEF1G CCL5 SLC12A2 ETS2 FCGBP TFF3 XBP1 MUC5AC MLPH ANXA2 CTSB CD55 S100A4 EPCAM FTH1 EEF1G CCL5 UGT2B17 ATP5F1A FTH1 SLC26A3 SLC26A2 MALL LGALS3 HMGB1 TUBB LDHA HSPD1 HSPA8 EEF1G FTH1 HNRNPA1L2 HSPA8 H19 OLFM4 EEF1G HSPA8 EIF4A1 FBL CXCL3 CXCL1 CXCL2 AREG OLFM4 CXCL14 CTSB BMP4 APCDD1 SERPINE2
Fraction of cells in group (%)
20 40 60 80 100
Mean expression in group
0.0 0.5 1.0
C
-log10(p_adj)
Dist. N vs Adj. N
Dist. N vs AD
Dist. N vs CA
Adj. N vs AD
Adj. N vs CA
AD vs CA
Comparison
D
Credible
Not credible
Comparison

Supplementary Figure 11. Characterization of Xenium epithelial clusters across patients and tissue types.

(A) UMAP visualization of subclustered epithelial cells from the Xenium Prime 5K TMA cohort (101 cores from 16 patients), colored by cluster, alongside stacked bar plots showing epithelial cluster composition across tissue types within each patient.
(B) Dot plot showing the top five marker genes for each epithelial cluster in (A). Dot size represents the percentage of cells expressing each gene, and color represents mean expression scaled within each gene.
(C) Heatmap of Mann-Whitney U test results comparing epithelial cluster composition across tissue types. Cell counts were aggregated at the patient-by-tissue-type level. Benjamini-Hochberg-adjusted P values are displayed as -log10-transformed values; greater color intensity indicates stronger evidence of a difference. Rows denote epithelial clusters, and columns denote tissue-type comparisons. * adjusted P < 0.05; ** adjusted P < 0.01; *** adjusted P < 0.001.
(D) Dot plot showing scCODA credibility results for changes in epithelial cluster composition across all tissue-type comparisons. Filled dots indicate credible changes, and dot color identifies the epithelial cluster. Cell counts were aggregated at the patient-by-tissue-type level; credibility was defined using the default inclusion probability threshold of 0.95. AD, conventional adenoma; CA, carcinoma; Adj. N, adjacent normal; Dist. N, distant normal.

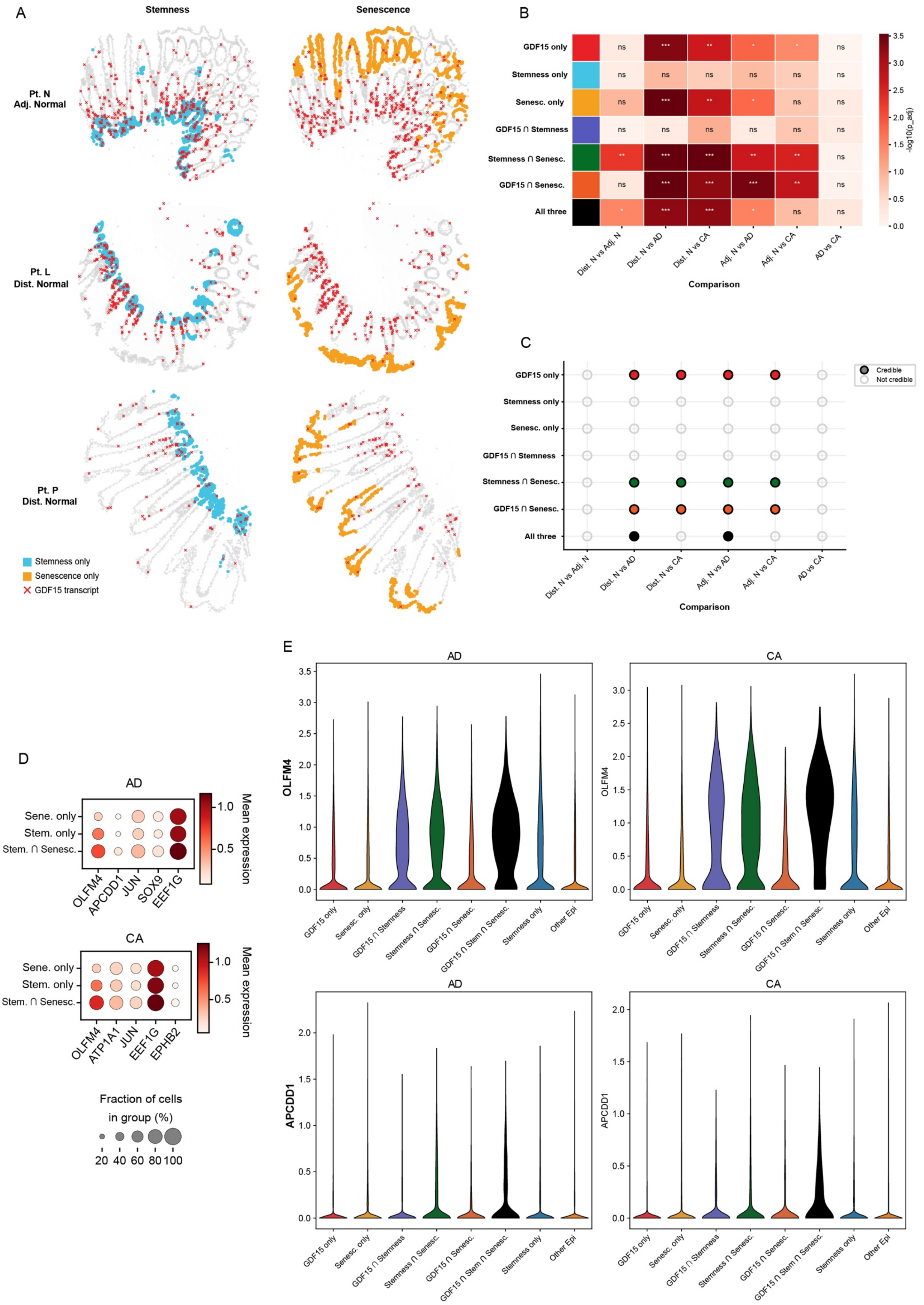
A
Stemness
Senescence
Pt. N
Adj. Normal
Pt. L
Dist. Normal
Pt. P
Dist. Normal
Stemness only
Senescence only
GDF15 transcript
B
GDF15 only
Stemness only
Senesc. only
GDF15 ∩ Stemness
Stemness ∩ Senesc.
GDF15 ∩ Senesc.
All three
Dist. N vs Adj. N
Dist. N vs AD
Dist. N vs CA
Adj. N vs AD
Adj. N vs CA
AD vs CA
Comparison
-log10(p_adj)
C
Credible
Not credible
Comparison
D
AD
Sene. only
Stem. only
Stem. ∩ Senesc.
OLFM4
APCDD1
JUN
SOX9
EEF1G
Mean expression
CA
ATP1A1
EPHB2
Fraction of cells
in group (%)
20 40 60 80 100
E
AD
CA
OLFM4
APCDD1
GDF15 only
Senesc. only
GDF15 ∩ Stemness
Stemness ∩ Senesc.
GDF15 ∩ Senesc.
GDF15 ∩ Stem ∩ Senesc.
Stemness only
Other Epi

Supplementary Figure 12. Spatial patterns and markers of stemness, senescence, and GDF15 hotspots across tissue types.

(A) Spatial maps showing stemness-only hotspots (blue) and senescence-only hotspots (yellow) in three representative normal cores. GDF15 transcripts are marked by red X symbols.

(B) Heatmap of Mann-Whitney U test results comparing the composition of stemness, senescence, and GDF15 hotspot categories across tissue types. Cell counts were aggregated at the patient-by-tissue-type level. Benjamini-Hochberg-adjusted P values are displayed as -log10-transformed values; greater color intensity indicates stronger evidence of a difference. Rows denote hotspot categories, and columns denote tissue-type comparisons. * adjusted P < 0.05; ** adjusted P < 0.01; *** adjusted P < 0.001.

(C) Dot plot showing scCODA credibility results for changes in stemness, senescence, and GDF15 hotspot composition across all tissue-type comparisons. Filled dots indicate credible changes, and dot color identifies the hotspot category. Cell counts were aggregated at the patient-by-tissue-type level; credibility was defined using the default inclusion probability threshold of 0.95. Adj. N, adjacent normal; Dist. N, distant normal.

(D) Dot plots showing the top five marker genes distinguishing Stemness ∩ Senescence hotspots from Stemness-only and Senescence-only hotspots in conventional adenoma (AD; top) and carcinoma (CA; bottom) cells. Dot size represents the percentage of cells expressing each gene, and color represents mean expression.

(E) Violin plots comparing mean OLFM4 (top) and APCDD1 (bottom) expression across local indicators of spatial association (LISA) hotspot categories in AD (left) and CA (right) tissues.

**Supplementary Table 1.** Clinical and histopathological features of the Visium adenoma cohort.

| ID | Gender | Age | Polyp Detail | Histology | Size |
|---|---|---|---|---|---|
| Pilot_large_1 | M | 67 | Right Colon | TVA | >10 mm |
| Pilot_small_1 | M | 54 | Sigmoid | TA | <5 mm |
| Pilot_large_2 | M | 63 | Sigmoid | TVA | >10 mm |
| R2_large_2 | M | 71 | Ascending Colon | TA | >10 mm |
| R2_small_2 | F | 73 | Left Colon | TA | <5 mm |
| R2_small_1 | M | 50 | Rectal | TA | <5 mm |
| R2_large_1 | M | 74 | Cecal | TVA | >10 mm |
| R3_large_1 | M | 56 | Transverse | TA | >10 mm |
| R3_large_7 | M | 66 | Colon | TA | >10 mm |
| R3_large_2 | F | 60 | Right Colon | TA | >10 mm |
| R3_large_6 | F | 60 | Left Colon | TA | >10 mm |
| R3_large_3 | M | 58 | Right Colon | TA | >10 mm |
| R3_large_5 | F | 62 | Ascending Colon | TA | >10 mm |
| R3_small_1 | M | 67 | Right Colon | TA | <5 mm |
| R3_large_4 | F | 69 | Ascending Colon | TA | >10 mm |
| R4_small_1 | M | 66 | Cecal | TA | <5 mm |
| R4_large_1 | M | 58 | Cecal | TA | >10 mm |
| R4_large_2 | M | 63 | Sigmoid | TVA | >10 mm |
| R4_large_3 | M | 58 | Sigmoid | TA/TVA | >10 mm |
| R4_large_5 | M | 52 | Sigmoid | TA | >10 mm |
| R4_small_2 | M | 64 | Cecal | TA | <5 mm |
| R4_large_4 | F | 70 | Cecal | TA | >10 mm |

**Supplementary Table 2.** Clinical and histopathological features of the Xenium TMA cohort.

| ID | Gender | Age | Location | # Cores | # sep_N | # N | #AD | #CA | # Mixed | Mixed Core Composition |
|---|---|---|---|---|---|---|---|---|---|---|
| J | F | 65 | Sigmoid | 5 | 0 | 2 | 1 | 1 | 1 | AD/CA |
| K | F | 73 | Right Colon | 6 | 2 | 0 | 1 | 3 | 0 | |
| L | M | 71 | Sigmoid | 7 | 2 | 0 | 3 | 1 | 1 | N/AD |
| M | M | 64 | Sigmoid | 4 | 0 | 2 | 0 | 1 | 1 | N/AD |
| N | M | 53 | Rectosigmoid | 8 | 2 | 2 | 2 | 2 | 0 | |
| O | M | 49 | Left Colon | 6 | 2 | 2 | 2 | 0 | 0 | |
| P | M | 59 | Sigmoid | 4 | 4 | 0 | 0 | 0 | 0 | |
| Q | F | 60 | Rectal | 7 | 0 | 1 | 6 | 0 | 0 | |
| R | F | 65 | Rectal | 6 | 2 | 0 | 2 | 1 | 1 | AD/CA |
| S | F | 67 | Right Colon | 7 | 2 | 2 | 2 | 1 | 0 | |
| T | M | 73 | Right Colon | 7 | 2 | 1 | 1 | 2 | 1 | N/AD |
| U | F | 42 | Rectal | 2 | 2 | 0 | 0 | 0 | 0 | |
| V | M | 81 | Right Colon | 6 | 2 | 0 | 2 | 2 | 0 | |
| W | F | 63 | Right Colon | 7 | 2 | 2 | 1 | 2 | 0 | |
| X | M | 60 | Right Colon | 8 | 2 | 2 | 2 | 2 | 0 | |
| Y | M | 71 | Right Colon | 11 | 0 | 4 | 2 | 2 | 3 | N/AD |
